# The Tuberous Sclerosis gene, *Tsc1*, represses parvalbumin+/fast-spiking properties in somatostatin-lineage cortical interneurons

**DOI:** 10.1101/699892

**Authors:** Ruchi Malik, Emily Ling-Lin Pai, Anna N Rubin, April M Stafford, Kartik Angara, Petros Minasi, John L Rubenstein, Vikaas S Sohal, Daniel Vogt

## Abstract

Medial ganglionic eminence (MGE)-derived somatostatin (SST)+ and parvalbumin (PV)+ cortical interneurons (CINs), have characteristic molecular, anatomical and physiological properties. However, mechanisms regulating their diversity remain poorly understood. Here, we show that conditional loss of the Tuberous Sclerosis (TS) gene, *Tsc1*, which inhibits mammalian target of rapamycin (MTOR), causes a subset of SST+ CINs, to express PV and adopt fast-spiking (FS) properties, characteristic of PV+ CINs. These changes also occur when only one allele of *Tsc1* is deleted, making these findings relevant to individuals with TS. Notably, treatment with rapamycin, which inhibits MTOR, reverses these changes in adult mice. These data reveal novel functions of MTOR signaling in regulating PV expression and FS properties, which may contribute to some neuropsychiatric symptoms observed in TS. Moreover, they suggest that CINs can exhibit properties intermediate between those classically associated with PV+ or SST+ CINs, which may be dynamically regulated by the MTOR signaling.

## Introduction

Tuberous Sclerosis (TS) is a disorder that affects multiple organ systems. Roughly half of those diagnosed with TS have autism spectrum disorder (ASD) or intellectual disability (ID), and ∼90% exhibit seizures ^1–4^. TS is caused by mutations in the *TSC1* and *TSC2* genes, which encode the HAMARTIN and TUBERIN proteins, respectively ^5,6^. HAMARTIN and TUBERIN proteins dimerize to form the Tuberous Sclerosis (TS) complex which inhibits the activity of the mammalian target of rapamycin (MTOR), a protein complex that is a rheostat for energy homeostasis and is also an important regulator of protein translation ^7^. Multiple proteins in the TS-complex/MTOR signaling pathway are either high confidence ASD-causative genes or underlie disorders with high ASD coincidence ^8,9^. This has relevance to the high rate of ASD in TS and potentially other TS-Associated Neuropsychiatric Disorders (TANDs), which are common in the syndrome ^10^. Uncovering how this pathway regulates neuronal development and function is therefore fundamental to understanding the molecular and cellular underpinnings of ASD and complex neuropsychiatric symptoms in TS.

Accumulating evidence suggests that neuropsychiatric disorders, such as ASD, and associated comorbidities like epilepsy, may be partially caused by changes in cortical GABAergic interneuron (CIN) function and connectivity, which leads to excitation/inhibition (E/I) imbalance in cortical circuits ^11–13^. While the role of MTOR signaling and *Tsc* genes on excitatory neurons has been studied for some time, relatively little is known about their roles in CIN development and function ^14–16^. CINs are the major source of cortical inhibition and are largely derived from the medial and caudal ganglionic eminences (MGE and CGE) ^17,18^. PV+ and SST+ CINs are derived from MGE and constitute ∼70% of all CINs. These cells have substantially different morphological and physiological properties ^19,20^. PV+ CINs exhibit fast-spiking (FS) firing properties and synapse onto soma/axons of excitatory neurons. By contrast, SST+ CINs have regular-spiking (RS) firing properties and target the distal dendrites of excitatory neurons ^20,21^.

Progenitor cells in the MGE that express the *Lhx6* transcription factor (TF) generate descendants that become either PV or SST CINs ^22–25^. Most studies investigating MGE-derived CIN fate and function have largely focused on the role of TFs ^23,24,26,27^, yet little is known about how cellular signaling influences CIN development. Recent work from us and others highlighted the importance of *Pten*, another inhibitor of MTOR signaling that acts upstream of *Tsc1*, on establishing proper numbers of MGE-derived CINs in the cortex ^28,29^. Therefore, we speculate that aberrant MTOR signaling in TS might lead to abnormalities in the development and function of MGE-derived CINs.

To investigate this, we conditionally deleted *Tsc1* in MGE-derived SST-lineage CINs, which allowed us to assess the impact of *Tsc1* loss/MTOR activity starting during early post-mitotic stages. We then investigated the role of *Tsc1*/MTOR signaling in CIN development, cell fate and physiology. Surprisingly, homozygous loss of *Tsc1* caused SST-lineage CINs to aberrantly exhibit cell fate and physiological properties of PV+/FS CINs. Notably, a subset of SST-lineage CINs resemble PV+/FS CINs in mice both homozygous and hemizygous for *Tsc1*, suggesting that these observations are relevant to humans with TS. In addition, this phenotype can be rescued by inhibiting MTOR during adult stages, suggesting that drugs currently being studied to treat TS, including rapamycin derivatives, may be effective in treating TS symptoms caused by CIN dysfunction. Overall, our findings demonstrate novel roles for *Tsc1* in the development and function of CINs. Notably, we propose that the choice between SST+ and PV+ cell fate and function is mediated in part by non-transcriptional processes, including cellular signaling events, suggesting a new avenue towards understanding these important cell types.

## Results

### Loss of *Tsc1* causes ectopic expression of PV in SST-lineage CINs

To test whether loss of *Tsc1* in SST-expressing post-mitotic CINs alters their development, we crossed *Tsc1*^*floxed*^ mice ^30^ and *SST-IRES-Cre* mice ^31^ (*SST-Cre* hereafter). The *Cre*-dependent tdTomato reporter, *Ai14* ^32^, was included to track the *Tsc1* WT, conditional heterozygous (cHet) and knockout (cKO) cells. *Tsc1* transcript was absent from *SST*-Cre; *Tsc1* cKO CINs in the neocortex at postnatal day (P) 35 (Supplementary Figs. 1a-d). At the same age, *SST-Cre*-lineage cHet and cKO cells in the neocortex had elevated levels of ribosomal subunit S6 phosphorylated at Serines 240 and 244 (Supplementary Figs. 1e-g), indicating increased MTOR activity.*Tsc1* cKOs had normal numbers of *SST-Cre*-lineage CINs (tdTomato+) in the neocortex but the cells had increased soma size (Figs. 1a-h).

**Figure 1:**
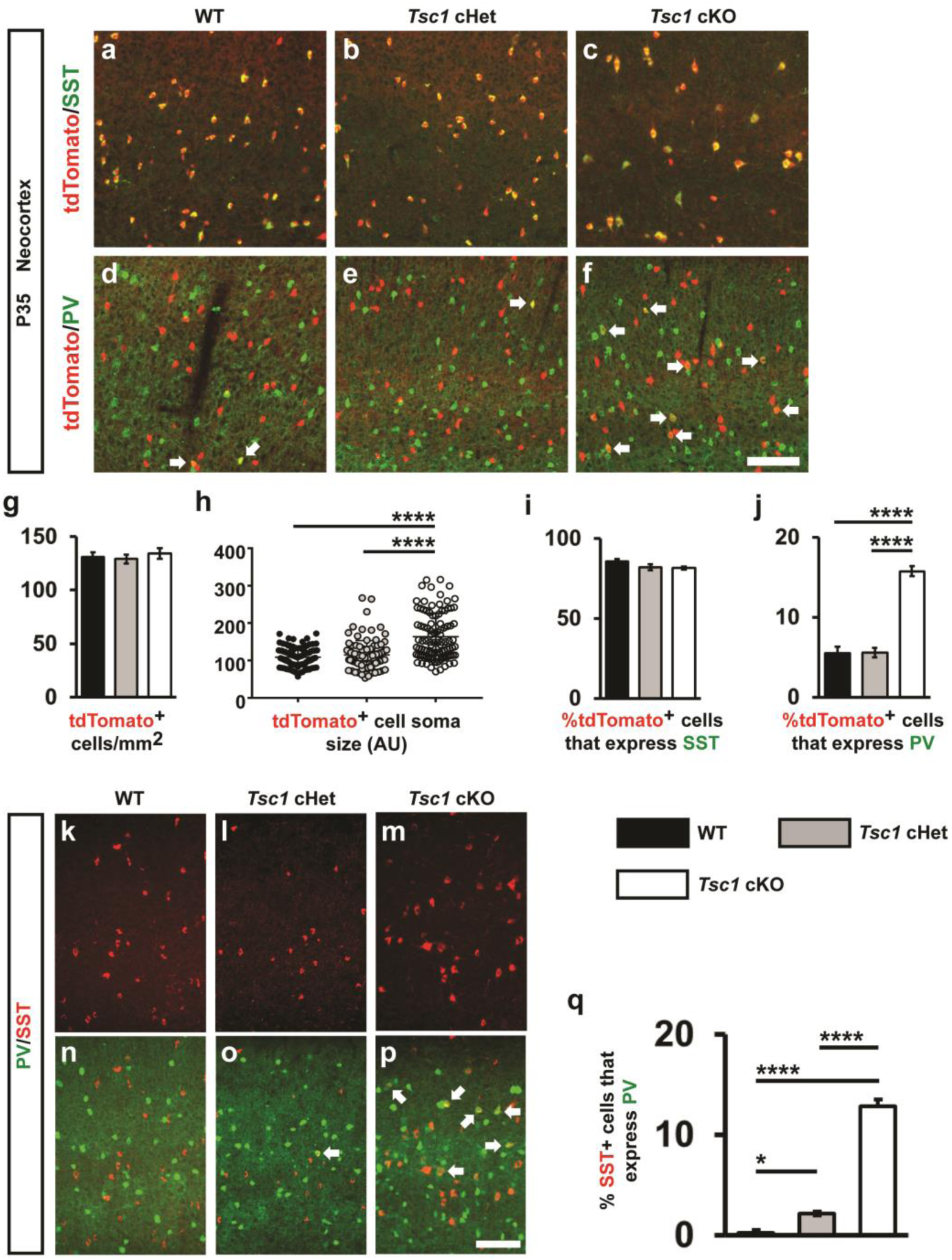
Post-mitotic deletion of *Tsc1* in *SST-Cre*-lineages results in increased PV expression. Coronal immuno-fluorescent images from WT, *Tsc1* cHets or *Tsc1* cKOs P35 neocortices, showing co-localization of tdTomato (*SST-Cre*-lineages) with either SST **(a-c)** or PV **(d-f)**. Arrows **(d-f)** denote cells in the *SST-Cre*-lineage that express PV. **(g)** Quantification of the number of tdTomato+ cells/mm^2^ (cell density) in the neocortex: n = 6 (WT), 6 (cHet) and 4 (cKO) mice, One-way ANOVA (F_2, 13_ = 0.35, P = 0.7). **(h)** Quantification of tdTomato+ cell soma size: n = 4 mice per group (25 cells measured from group), (F_2, 297_ = 44.9, P < 0.0001; cKO vs. WT and cHet, p < 0.0001). (AU) arbitrary units. **(g, h)** Quantification of the %tdTomato+ cells that co-express either SST **(i)** or PV **(j)**: Chi-squared test with Yate’s correction, (p < 0.0001, all groups), n = 4. **(k-p)** Immuno-fluorescent images showing SST and PV co-labeling in the neocortex. **(q)** Quantification of the %SST+ CINs that co-express PV: Chi-squared test with Yate’s correction, (WT vs cHet, p = 0.03, cKO vs. WT or cHet, p < 0.0001), n = 3 mice, all groups. Data are represented as mean ± SD in **(h)** and ± SEM in all other graphs. Scale bars in **(f, p)** = 100µm. * p < 0.05, **** p < 0.0001.

SST-lineage CINs differentiate from MGE progenitors that also give rise to PV+ CINs. Thus, we investigated the expression of SST and PV at P35 in the neocortex. While the % of tdTomato+ CINs that expressed SST was unchanged, there was ∼3-fold increase in the % of *SST-Cre*-lineage CINs that expressed PV in cKO mice (Figs. 1a-f, i, j). These data suggest that SST and PV proteins could be co-expressed in a subset of *SST-Cre*-lineage CINs in cKO mice. To test this, quantified WT, *Tsc1* cHet and cKO neocortices for SST and PV co-labeled CINs at P35 in the neocortex. While there were almost no co-labeled CINs in WT, ∼2% and 13% co-labeled CINs were observed in cHets cKOs, respectively (Figs. 1k-q). A similar phenotype of ectopic PV expression in SST+ CINs after *Tsc1* deletion was observed in *Nkx2.1-Cre; Tsc1* conditional mutants (Fig. S2). In these crosses, *Tsc1* was conditionally deleted using *Nkx2.1-Cre*, which begins to express in MGE progenitors ^33^. Together, these data suggest that *Tsc1* represses PV expression in SST-lineages and in its absence, a subset of SST+ CINs develop a dual molecular identity.

### SST+ CINs in *Tsc1* cKOs exhibit fast-spiking physiological properties

Next, we assessed the physiological properties of SST-lineage CINs in the *Tsc1* cKOs. We obtained *ex-vivo* patch clamp recordings from layer 5 *SST-Cre*-lineage CINs (tdTomato+) in WT, cHet and cKO mice (example cells, Fig. 2a, and recordings, Fig. 2b). WT CINs had RS physiological properties, including strong spike frequency accommodation (SFA), wider spikes, relatively slow action potential (AP) rate of rise (max dV/dt) and relatively small fast afterhyperpolarization (fAHP) amplitudes (Fig. 2), as expected ^19,20,34^. Consistent with the increased soma size, known to be regulated by *Tsc1*/MTOR signaling ^34,35^, and the increase in cKO soma size and dendritic complexity (Supplementary Figs. 4a-c), loss of *Tsc1* decreased the input resistance (R_in_) and produced a corresponding increase in the rheobase (current threshold) of CINs in cHets and cKOs (Figs. 2c, f). A reduction in these properties predicts that *Tsc1* loss would lower the excitability of SST+ CINs. However, *Tsc1* cKO CINs had increased firing output (Figs. 2d, g–h). This increase was most pronounced within a subset of *SST-Cre* lineage CINs with FS electrophysiological properties (Fig. 2h and Supplementary Fig. 3). This shift from a RS to FS phenotype was most prevalent in CINs from cKO mice (Figs. 2g, h), and was consistent with the increased % of tdTomato+ CINs expressing PV (Fig. 1). Notably, most of the SST-lineage CINs classified as FS in the *Tsc1* cKOs expressed PV (supplementary Figs. 4d-f). While a small subset of CINs in cHet and cKO mice completely switched their physiology from RS to FS, ∼30%, the firing and single AP properties of most *SST-Cre*-lineage CINs in cHet and cKO mice shifted towards a FS-like physiology. Significant shifts were observed in AP threshold, AP rate of rise (max. dV/dt), AP duration and fAHP amplitudes (Figs. 2i–m). Together, these analyses showed that while loss of *Tsc1* reduced the excitability of SST+ CINs, their overall firing output was higher due to the increased proportion of SST+ CINs with PV-like/FS properties. Notably, most of the SST-lineage CINs classified as FS CINs in the *Tsc1* cKOs expressed PV (Supplementary Figs. 4a–c).

**Figure 2:**
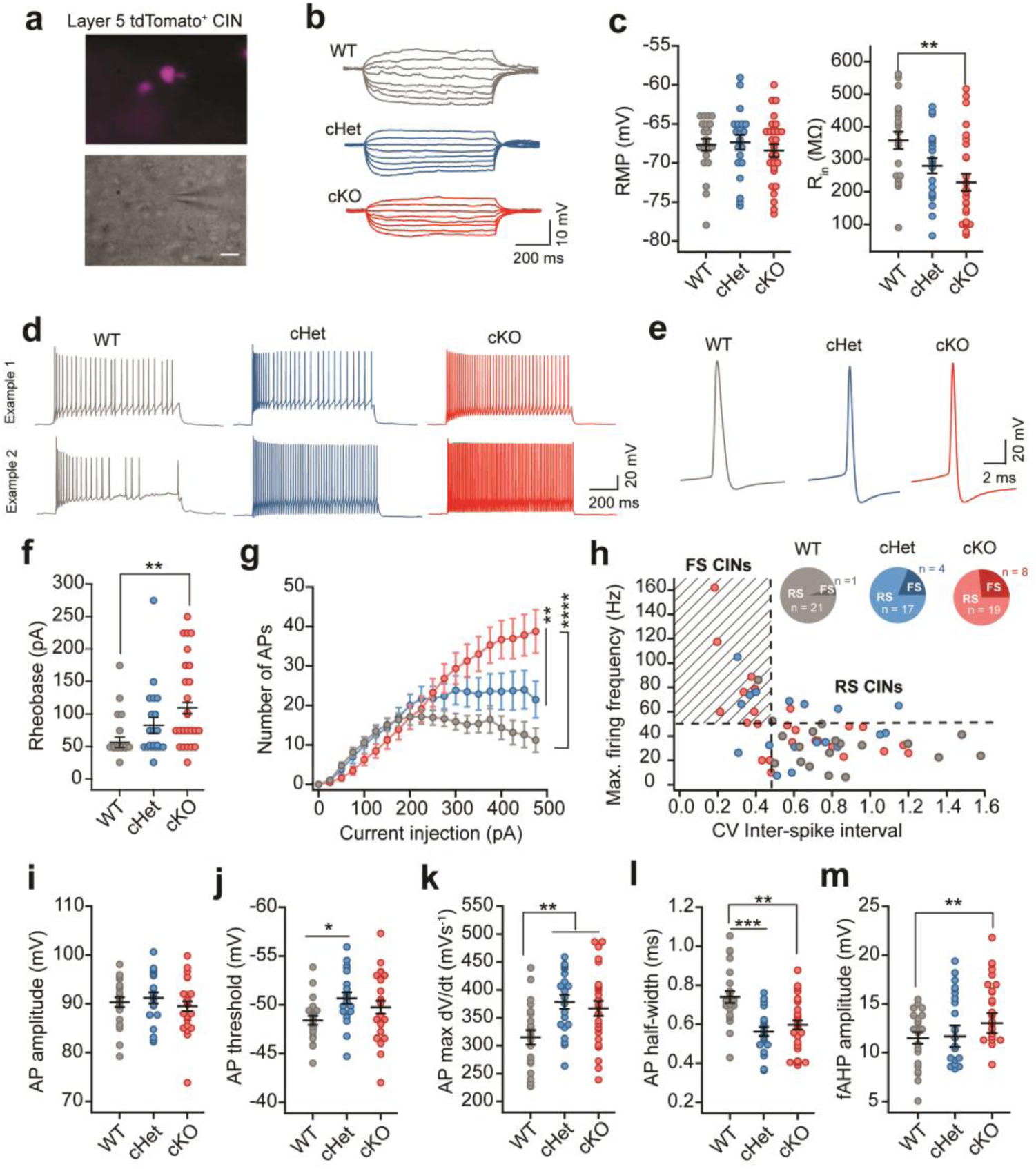
*SST-Cre*-lineage CINs adopt fast-spiking (PV-like) electrophysiological properties after deletion of *Tsc1*. **(a)** Fluorescence (top) and DIC (bottom) images illustrating that patch clamp recordings were obtained from layer 5 tdTomato+ *SST-Cre* lineage CINs. Scale bar is 20 µm. **(b)** Example voltage traces in response to subthreshold hyperpolarizing and depolarizing current injections in SST+ CINs from WT (grey), *Tsc1* cHet (blue) and *Tsc1* cKO mice (red). **(c)** Comparison of resting membrane potential (RMP), (F_2, 67_ = 0.41, P = 0.66) and input resistance (R_in_), (F_2, 67_ = 6.49, P = 0.002; WT vs. cKO p = 0.001), One-way ANOVA with Tukey’s post-test. **(d)** Example voltage traces in response to suprathreshold current injections from SST-lineage CINs in WT, cHet and cKO mice. Example traces from two different cells are shown to illustrate the diversity in firing pattern of cHet and cKO SST-lineage CINs. **(e)** Example voltage traces showing single action potentials (APs) in WT, cHet and cKO CINs. **(f)** SST-lineage CINs in cKO mice have significantly higher rheobase: One-way ANOVA with Tukey’s post-test (F_2, 67_ = 5.06, P = 0.009; WT vs. cKO, p = 0.006). **(g)** Number of APs fired in response to steps of depolarizing current injections is plotted. Firing output is significantly higher for SST+ CINs in cKO and cHet groups as compared to WT group: Two-way ANOVA with Tukey’s post-test, Genotype: (F_2, 1320_ = 26.8, P < 0.0001); WT vs. cHet, and WT vs. cKO, p < 0.0001; cHet vs. cKO, p = 0.008). **(h)** Maximum firing frequency of SST+ CINs from WT, cHet and cKO mice is plotted against the coefficient of variance (CV) of inter-spike interval. Inset, pie charts showing the proportion of SST-lineage CINs with regular-spiking (RS) and fast-spiking (FS) physiological properties in WTs, cHets and cKOs. (**i–m)** Comparison of AP amplitude **(i):** (F_2, 67_ = 0.67, P = 0.51), AP threshold **(j):** (F_2, 67_ = 3.48, P = 0.03; WT vs. cHet p = 0.03), AP max dV/dt **(k)**: (F_2, 67_ = 6.39, P = 0.002; WT vs. cHet p = 0.004, WT vs. cKO p = 0.009), AP half-width **(l)** (F_2, 67_ = 11.04, P < 0.0001; WT vs. cHet p = 0.0001, WT vs. cKO p = 0.001); and fAHP amplitude **(m)**: (F_2, 67_ = 5.27, P = 0.007; WT vs. cKO p = 0.005) of SST+ CINs recorded from WT, cHet and cKO mice; One-way ANOVA with Tukey’s post-test. WT, n = 22 cells from 4 mice; cHet, n = 21 cells from 3 mice; cKO, n = 27 cells from 4 mice; data are presented as mean ± S.E.M. * p < 0.05; ** p < 0.01, *** p < 0.001, **** p < 0.0001.

### Increased Kv3.1 expression underlies the switch in physiology of the *SST+* CINs in *Tsc1* cKOs

Since *Tsc1* deletion resulted in a shift towards FS physiology within *SST-Cre*-lineages, we asked whether *Tsc1* cHets and cKOs had increased levels of the fast-inactivating Kv3.1 potassium channel. Kv3.1 expression is primarily restricted to PV+ CINs and contributes to many of their fast-spiking properties (like short AP half-width and large fAHP) ^36,37^ (Supplementary Fig. 5). Indeed, we found that increased proportions of *SST-Cre*-lineage CINs co-expressed Kv3.1 in cHets and cKOs (Figs. 3a, b).

**Figure 3:**
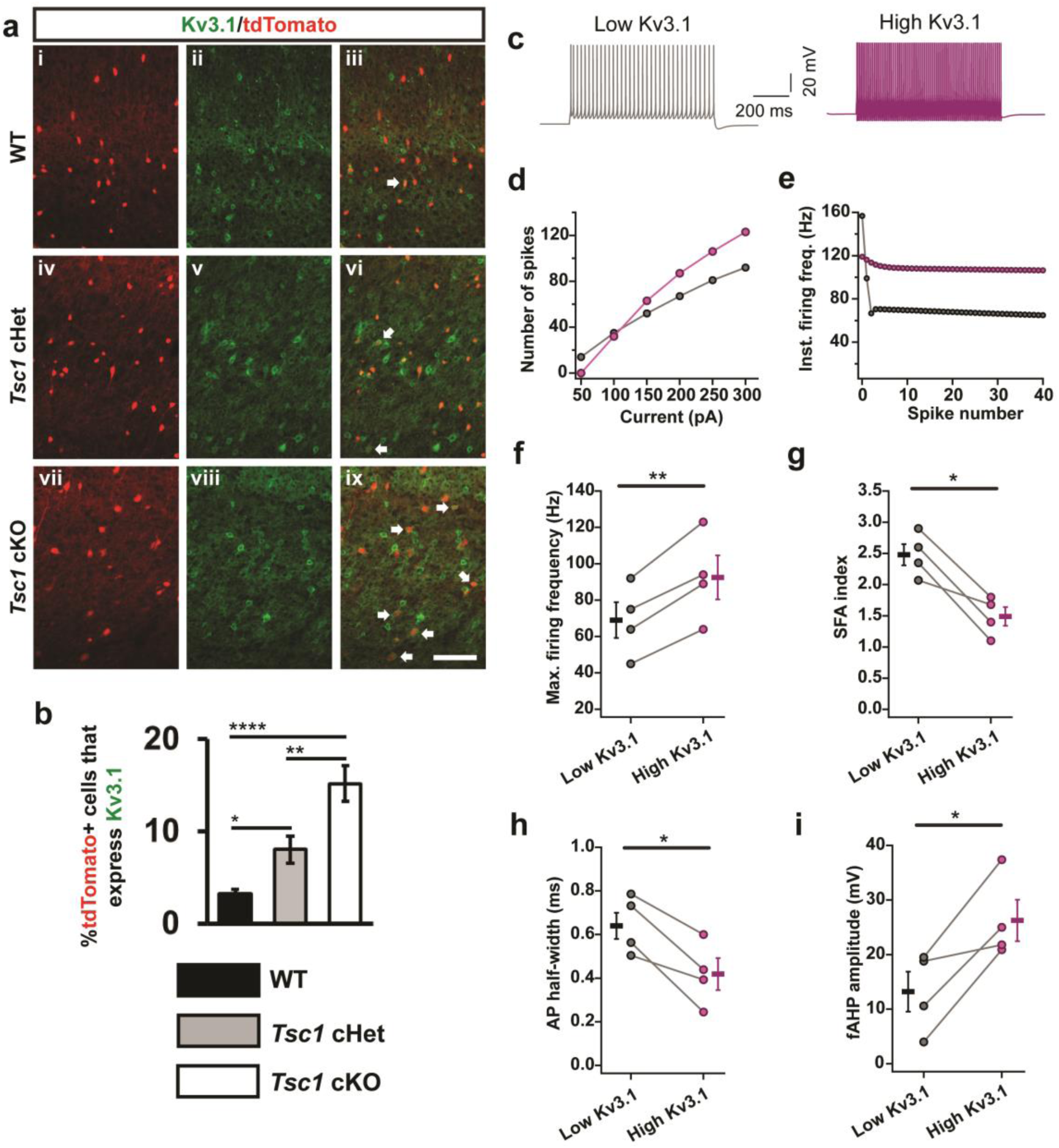
Increased Kv3.1 channel expression in *SST-Cre*-lineage CINs underlies the fast-spiking electrophysiological properties. **(a)** P35 coronal sections showing immuno-fluorescent labeling of Kv3.1 and tdTomato+ cells from *SST-Cre*-lineages in the neocortices of WT (i–iii), *Tsc1* cHets (iv–vi) and cKOs (vii–ix). Arrows denote co-labelled cells. Scale bar is 100 µm. **(b)** Quantification of the %tdTomato+ cells expressing Kv3.1: Chi-squared test with Yate’s correction, WT vs. cHet, p = 0.02 or cKO, p < 0.0001, and cHet vs. cKO, p = 0.008, n = 3 (all groups). **(c)** Example firing traces from a model of a layer 5 SST+ CIN with low expression of Kv3.1 (grey) and high expression of Kv3.1 (purple). (**d-e**) Firing data from one representative model is shown. Increasing the conductance of Kv3.1 channel increases the number of APs fired in response to depolarizing current injections **(d)** and decreases the change in instantaneous firing frequency during a train of APs **(e)**. Increasing the Kv3.1 conductance in four different models of SST+ CINs increases the maximum firing frequency: paired two-tailed t test (t_3_ = 8.1, p = 0.003) **(f)** and decreases the spike frequency accommodation: SFA; paired two-tailed t test (t_3_ = 4.9, p = 0.016) **(g)**. Increasing Kv3.1 conductance in models of SST+ CINs decreases the duration of APs: paired two-tailed t test (t_3_ = 4.7, p = 0.018) **(h)** and increases the fAHP: paired two-tailed t test (t_3_ = 3.8, p = 0.03) **(i)**. Data are presented as mean ± S.E.M. * p < 0.05, ** p < 0.01, **** p < 0.0001.

To understand the effects of increased Kv3.1 channel expression on the physiological properties of SST+ CINs, we used biophysical simulations of four different models of neocortical layer 5 SST+ CINs ^38^. Specifically, we asked if an increase in somatic Kv3.1 conductance density was sufficient to produce a transition from RS to FS properties (similar to the switch observed in *Tsc1* cKOs). In all four models, increasing Kv3.1 conductance density increased the firing output (average increase of 28%), reduced spike frequency accommodation (40% reduction), reduced AP half-width (32% reduction) and increased fAHP amplitude (80% increase) (Figs. 3c–i and Supplementary Fig. 6). These results suggest that *Tsc1* inhibits FS properties via its regulation of Kv3.1 in SST-lineage CINs.

### Reduced inhibitory synaptic output of SST+ CINs in *Tsc1* cKOs

Given that loss of *Tsc1* affected the excitability and firing output of SST+ CINs, we wondered how these changes might affect inhibitory synaptic output onto nearby pyramidal neurons. Thus, we compared spontaneous inhibitory postsynaptic currents (sIPSCs) recorded from cortical layer 5 pyramidal neurons in WT, cHet and cKO mice (Figs. 4a). Surprisingly, the frequency and amplitude of sIPSCs was lower in cHet and cKO mice (Fig. 4b). We had expected that the increase in firing output might result in increased spontaneous inhibitory transmission. This reduction in spontaneous inhibitory output could be caused by a reduction in numbers of inhibitory synapses and/or probability of release at these synapses. Towards elucidating the respective contributions of these various possibilities, we added TTX (1µM) to the recording solution, to remove large amplitude, AP-dependent IPSCs (and therefore any effects of changes in excitability). Similar to sIPSCs, the frequency and amplitude of miniature IPSCs (mIPSCs) were lower in *Tsc1* cKOs (Figs. 4c, d). These findings suggested that the loss of *Tsc1* reduced the strength of inhibitory synaptic transmission by altering the number and/or release probability at inhibitory synapses.

**Figure 4:**
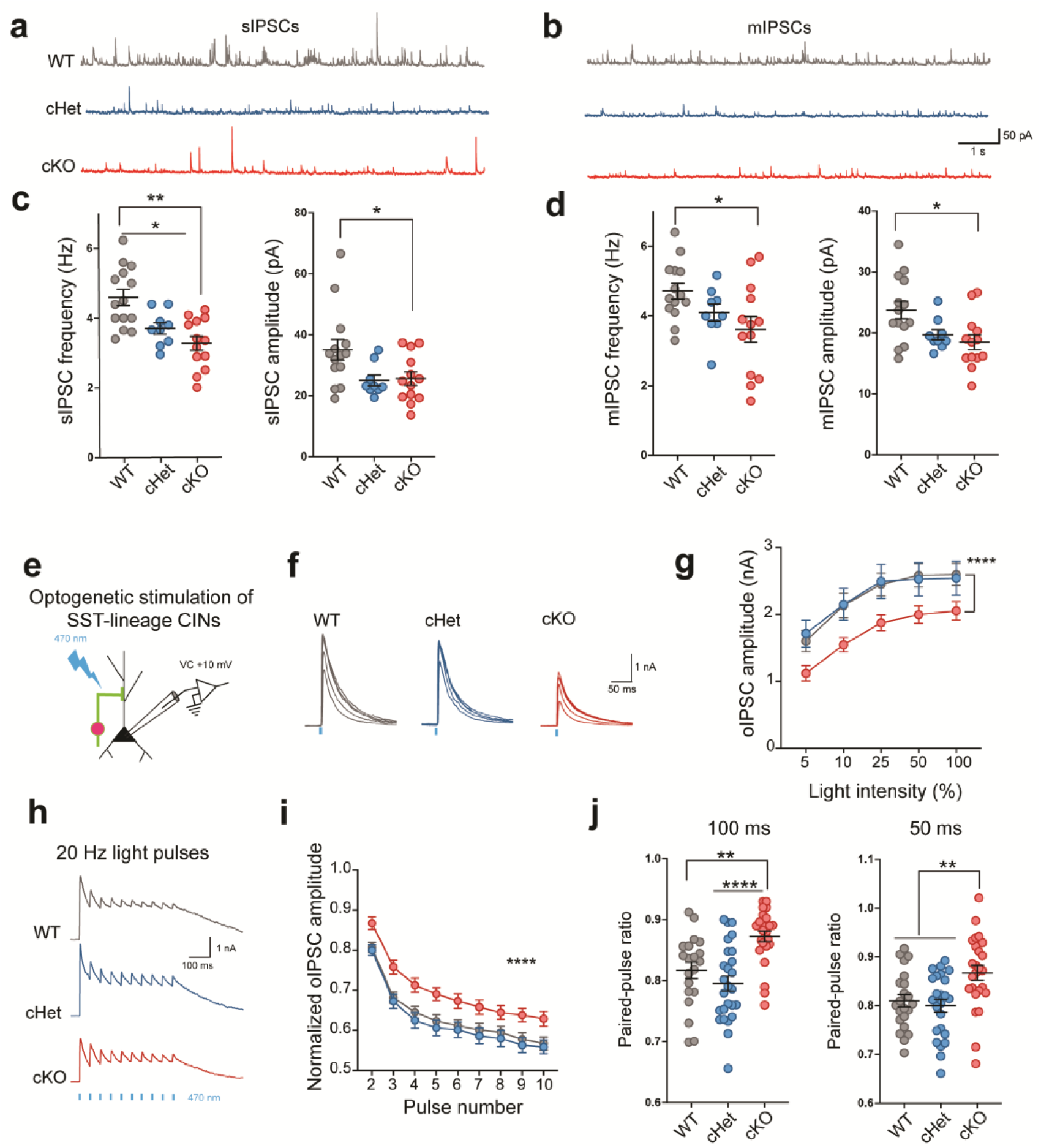
Reduced inhibitory synaptic output of SST+ CINs in *Tsc1* cKOs. Example traces showing sIPSCs **(a)** and mIPSCs **(b)** recorded during voltage clamp recordings from layer 5 pyramidal neurons in WT (grey), cHet (blue) and cKO (red) mice. Comparison of the frequency and amplitude of sIPSCs **(c)** and mIPSCs **(d)** recorded from the three groups of mice (WT, n = 14 cells from 3 mice; cHet, n = 9 cells from 2 mice; cKO, n = 13 cells from 3 mice). One-way ANOVA with Tukey’s post-test, sIPSC Frequency: (F_2, 33_ = 11.16, P = 0.0002; WT vs cHet p = 0.02; WT vs cKO p = 0.0002); sIPSC Amplitude: (F_2, 33_ = 4.3, P = 0.02; WT vs cKO p = 0.03). mIPSC Frequency: (F_2, 33_ = 3.2, P = 0.04; WT vs cKO p = 0.04); mIPSC Amplitude: (F_2, 33_ = 5.1, P = 0.01; WT vs cKO p = 0.01). **(e)** Experimental design: voltage clamp recordings were obtained from layer 5 pyramidal neurons while activating ChR2-expressing SST-lineage CINs. **(f)** Example optogenetically-activated IPSCs (oIPSCs) recorded from pyramidal neurons in WTs (21 cells from 3 mice), cHets (24 cells from 3 mice) and cKOs (25 cells from 3 mice) in response to 5 ms blue light (470 nm) flashes of increasing light-intensity (4mW, maximum intensity). **(g)** Average oIPSC amplitudes evoked in response to single light pulses of increasing light intensity. The oIPSC amplitudes in cKOs are smaller in comparison to WTs and cHets: Two-way ANOVA with Tukey’s post-test (F_2, 335_ = 16.19, P < 0.0001; WT and cHet vs. cKO p < 0.0001). **(h)** Example oIPSCs recorded in response to 20 Hz light pulse trains in WTs, cHets and cKOs. **(i)** The attenuation in oIPSC amplitude during repeated activation of SST-lineage CINs by 20 Hz light pulse trains. The % attenuation is lower in cKOs compared to WTs and cHets: Two-way ANOVA with Tukey’s post-test (F_2,612_ = 56.6, P < 0.0001; WT and cHet vs. cKO p < 0.0001). **(j)** Paired-pulse ratio of oIPSCs in response to pairs of light pulses: One-way ANOVA with Tukey’s post-test, 100 ms inter-pulse interval: (F_2, 67_ = 12, P = 0.0002; WT vs cKO p = 0.002; cHet vs cKO p < 0.0001) and 50 ms inter-pulse interval: (F_2, 67_ = 7.4, P = 0.001; WT vs cKO p = 0.002; cHet vs cKO p = 0.008). Data are presented as mean ± S.E.M. * p < 0.05, ** p < 0.01, **** p < 0.0001.

While we found significant reductions in spontaneous inhibitory output to cortical pyramidal neurons, we could not estimate the specific contribution of SST+ CINs to this reduction by analyzing sIPSCs and mIPSCs. To directly address this, we expressed ChR2-eYFP in *SST-Cre* lineage CINs (WT, cHet and cKO). By delivering pulses of blue (470nm) light, while making voltage-clamp recordings, we measured optogenetically-evoked IPSCs (oIPSCs) from *SST*-lineage CINs onto layer 5 pyramidal neurons (Figs. 4e, f). In line with the observed reduction in spontaneous inhibitory output, oIPSCs were also significantly smaller in *Tsc1* cKOs (Fig. 4g). To measure the presynaptic release probability, we measured the frequency dependent attenuation of oIPSCs and paired-pulse ratio (PPR). The attenuation of oIPSCs in response to 20 Hz light pulses was significantly lower in cKOs (Figs. 4h, i). Furthermore, cKOs had significantly higher paired-pulse ratios (PPRs) (Fig. 4j). These results suggest that *Tsc1* deletion decreases the evoked inhibitory output at least in part by reducing presynaptic release probability in SST-lineage CINs.

The axons of PV+/FS CINs target the soma and axon initial segment of pyramidal neurons, while the axons of SST+ CINs target their distal dendrites ^20,21^. Thus, we tested whether *Tsc1* deletion caused aberrant axon targeting (proximal dendrite/soma vs. distal dendrite) in a subset of SST-lineage CINs. For this, we obtained paired patch clamp recordings from connected layer 5 pyramidal neurons (Pyr) and tdTomato+ *SST-Cre*-lineage CINs (Fig. 5a) and analyzed the unitary IPSCs (uIPSCs) in postsynaptic Pyr neurons evoked by action potentials in presynaptic SST+ CINs (Fig. 5b). Since uIPSCs generated in response to SST+ CINs arrive at the distal dendrites, the amplitude and kinetics of these currents are significantly different from IPSCs originating near the soma of pyramidal neurons ^39^. To validate this, we compared the properties of CIN→ Pyr uIPSCs in *PV-Cre*-lineage ^40^ CIN-pyramidal neuron vs. *SST-Cre*-lineage CIN-pyramidal neuron pairs. In our recordings from WT mice, all *PV-Cre* lineage CINs had FS physiology and all *SST-Cre* lineage CINs had RS physiology (Fig. 5c). A comparison of the kinetics of SST→Pyr uIPSCs and PV→ Pyr uIPSCs in WT mice showed that the uIPSC amplitude, rising slope and total charge were all significantly larger for PV→Pyr pairs than SST→Pyr pairs (Figs. 5d–f). Next, we recorded SST→Pyr uIPSCs in cKO mice. As expected, most *SST-Cre*-lineage CINs in cKOs had RS properties but a subset had FS properties (Fig. 5c). Strikingly, in our paired recordings from cKOs, the amplitude, kinetics and total charge uIPSCs did not differ for RS/SST→ Pyr vs. FS/SST→Pyr pairs (Figs. 5d–f). These results suggest that while *Tsc1* deletion caused a switch in physiological properties and molecular identity of *SST-Cre* lineage CINs towards a PV-like/FS physiology, it does not affect their axonal targeting. This finding together with our finding of lower presynaptic release probability explains why, despite the increased firing output and shift towards PV-like phenotypes, inhibitory output mediated by SST+ CINs is reduced in *Tsc1* cKOs.

**Figure 5:**
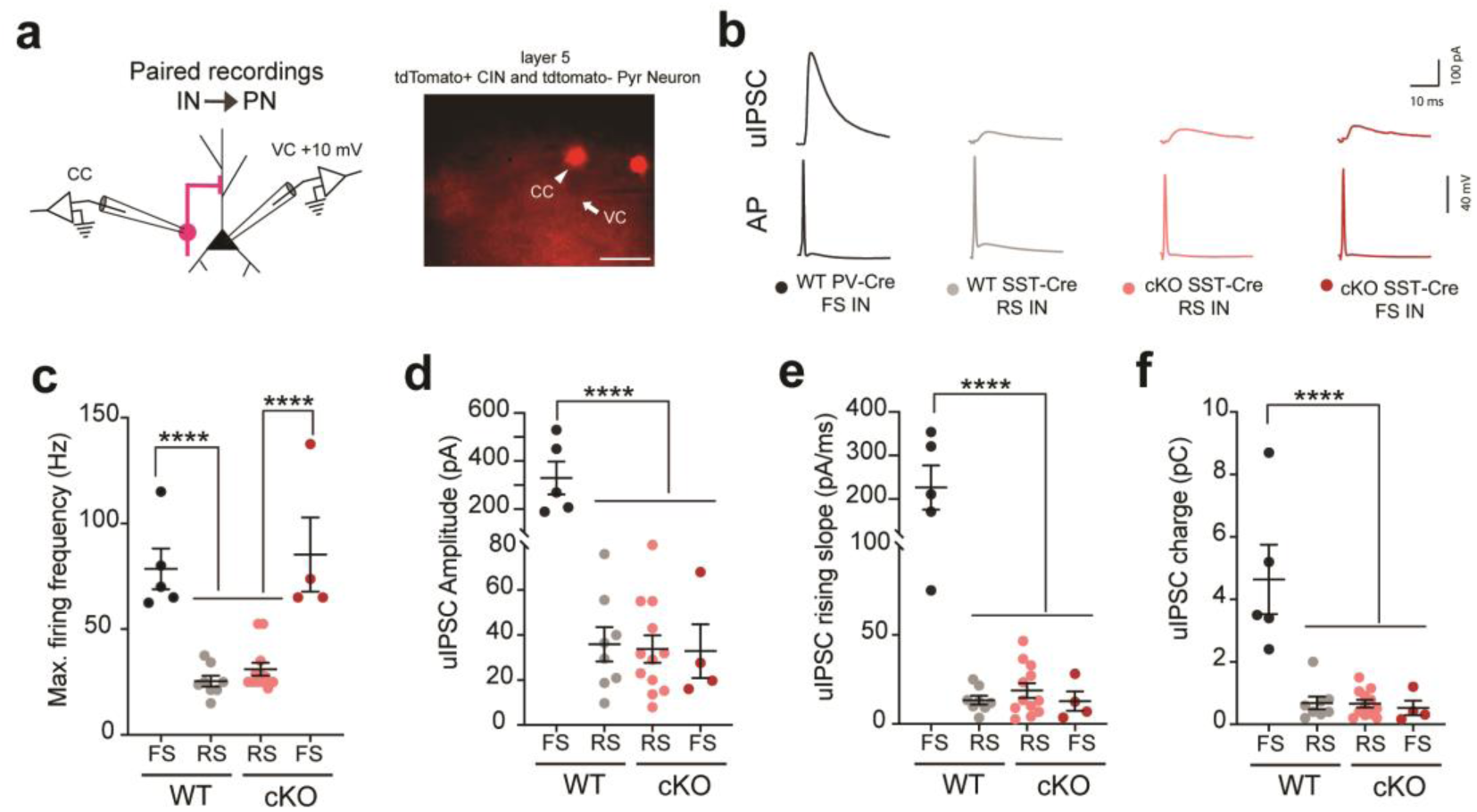
Fast spiking/*SST-Cre* CINs in *Tsc1* cKOs send their outputs to distal dendrites of pyramidal neurons. **(a)** Left, Experimental design: Simultaneous recordings were obtained from connected layer 5 tdTomato+ CINs, (current clamp, CC) and pyramidal neurons, (voltage clamp, VC), +10 mV. Right, representative image showing patch pipettes targeting tdTomato+ CIN (arrow head) and tdTomato-pyramidal neuron (arrow). Scale bar is 50µm. **(b)** Example unitary inhibitory post synaptic currents (uIPSCs) recorded in pyramidal neurons generated in response to action potentials (AP) in WT FS/*PV-Cre* CINs (black, 5 pairs), WT RS/*SST-Cre* CINs (grey, 8 pairs), cKO RS/*SST-Cre* CINs (salmon, 12 pairs) and cKO FS/*SST-Cre* CINs (red, 4 pairs). **(c)** Maximum firing frequency of four groups of CINs is compared: One-way ANOVA with Tukey’s post-test: (F_3, 25_ = 20, P < 0.0001; FS vs. RS p < 0.0001). **(d–f)** uIPSC amplitude, rising slope and total charge is compared. One-way ANOVA with Tukey’s post-test, uIPSC amplitude **(d)**: (F_3, 25_ = 29, P < 0.0001; WT FS/*PV-Cre* vs. all groups p < 0.0001); rising slope **(e)**: (F_3, 25_ = 28.11, P < 0.0001; WT FS/PV-Cre vs. all groups p < 0.0001); and total charge **(f)**: (F_3, 25_ = 18, P < 0.0001; WT FS/PV-Cre vs. all groups p < 0.0001). Data are presented as mean ± S.E.M. **** p < 0.0001.

### Cell autonomous role for *Tsc1* in regulating PV expression in CINs

Since *Tsc1* cKOs exhibited reduced inhibitory tone in the neocortex (Fig. 4), we asked if the increased PV expression in *SST-Cre*-lineages is cell autonomous, as PV expression can be modulated by circuit activity ^41,42^. We utilized an MGE transplantation assay that introduces MGE progenitors in small numbers to a WT cortex for *in vivo* maturation ^43^. MGE tissue from WT, *Tsc1*^*Floxed/+*^ or *Tsc1*^*Floxed/Floxed*^ was isolated from E13.5 embryos; dissociated cells were transduced with *DlxI12b-Cre* expressing lentiviruses (Fig. 6a). The *DlxI12b* enhancer biases expression to GABAergic neurons and has been used to express genes efficiently in developing and mature CINs ^22,28,44^. These MGE cells were then transplanted into WT neonatal cortex and allowed to develop. After 35 days post-transplant (DPT), we assessed the proportion of transplanted cells (tdTomato+) that expressed PV. *Tsc1* cKO cells had increased soma size and ∼2-fold increase in PV expression compared to WT (Figs. 6b-d, f, g). Finally, we expressed the human *TSC1* gene from the same lentiviral vector, to test if human *TSC1* could compliment the mouse *Tsc1* depleted CINs and found that it was able to rescue the increase in soma size and PV expression (Figs. 6e-g).

**Figure 6:**
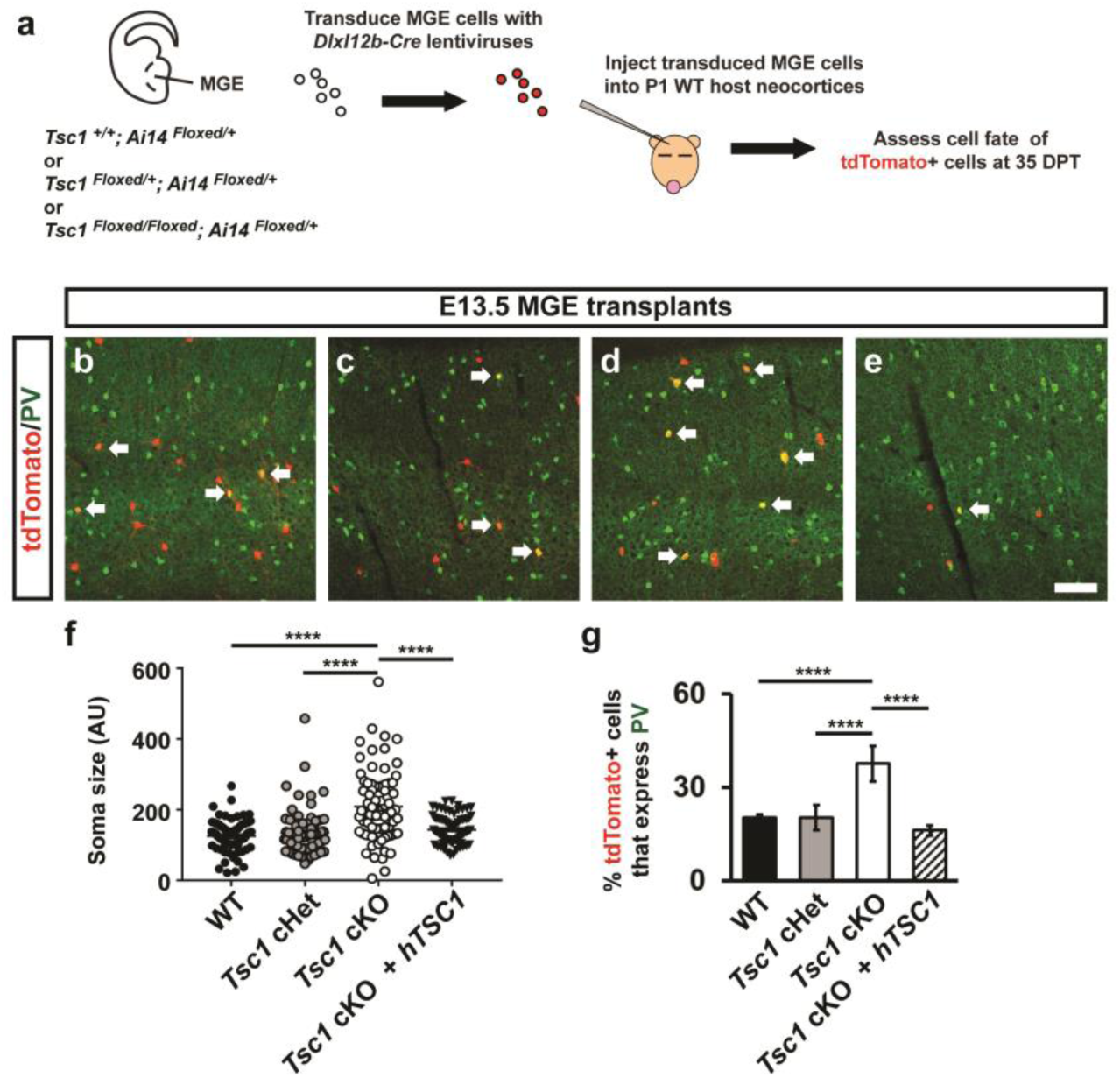
Cell autonomous increase in PV expression in *Tsc1* null MGE-derived CINs. **(a)** Schema depicting the transplantation procedure. E13.5 *Ai14*^*Floxed/+*^ MGE cells (tdTomato+) that were either *Tsc1*^*+/+*^, *Tsc1*^*Floxed/+*^ or *Tsc1*^*Floxed/Floxed*^ were transduced with a *DlxI12b-Cre* lentivirus, transplanted into WT P1 neocortices, then developed for 35 days post-transplant (DPT). Coronal immuno-fluorescent images of neocortices transplanted with *Tsc1*^*+/+*^ (WT), *Tsc1*^*Floxed/+*^ (cHet) or *Tsc1*^*Floxed/Floxed*^ (cKO) MGE cells show co-expression of tdTomato and PV **(b-d)**. In addition, human *TSC1* was expressed in *Tsc1*^*Floxed/Floxed*^ (cKO) MGE cells and assessed in the same manner **(e)**. Arrows point to co-expressing cells. Quantification of soma size of the tdTomato+ cells **(f):** One-way ANOVA, (F_3, 296_ = 24.1, P < 0.0001; cKO vs. all groups, p < 0.0001), n = 3 mice (25 cells counted from each). (AU) arbitrary units. Quantification of the proportion of tdTomato+ cells that express PV **(g)**: Chi-squared test with Yate’s correction, (cKO vs. all groups, p < 0.0001), n = 3, all groups. Data are represented as mean ± SD in **(f)** and ± SEM in **(g)**. Scale bar in **(e)** = 100µm. **** p < 0.0001.

### Five-day rapamycin treatment decreases PV expression and fast-spiking properties in *Tsc1* cKO *SST-Cre*-lineage CINs

Based on the preceding results, we hypothesized that *Tsc1* normally represses PV-like/FS properties in SST+ CINs by inhibiting MTOR activity. We thus administered the MTOR inhibitor, rapamycin, to test whether this treatment might reverse/rescue the effects of *Tsc1* loss in young adult (eight-week old) mice. *Tsc1*, cHets and cKOs were treated with rapamycin or vehicle starting at eight weeks of age, for five consecutive days (Fig. 7a). A day after the last dose, mice were assessed for soma size, PV expression and electrophysiology. To assess rapamycin’s efficacy, we probed for pS6 in tdTomato+ CINs. Vehicle-treated *Tsc1* cKO CINs had increased co-expression of pS6 compared to the cHet groups and rapamycin treatment significantly reduced these levels (Supplementary Figs. 7a-e). Notably, rapamycin treatment did not alter the cell density of tdTomato+ CINs or those expressing SST (Supplementary Figs. 7f, g).

**Figure 7:**
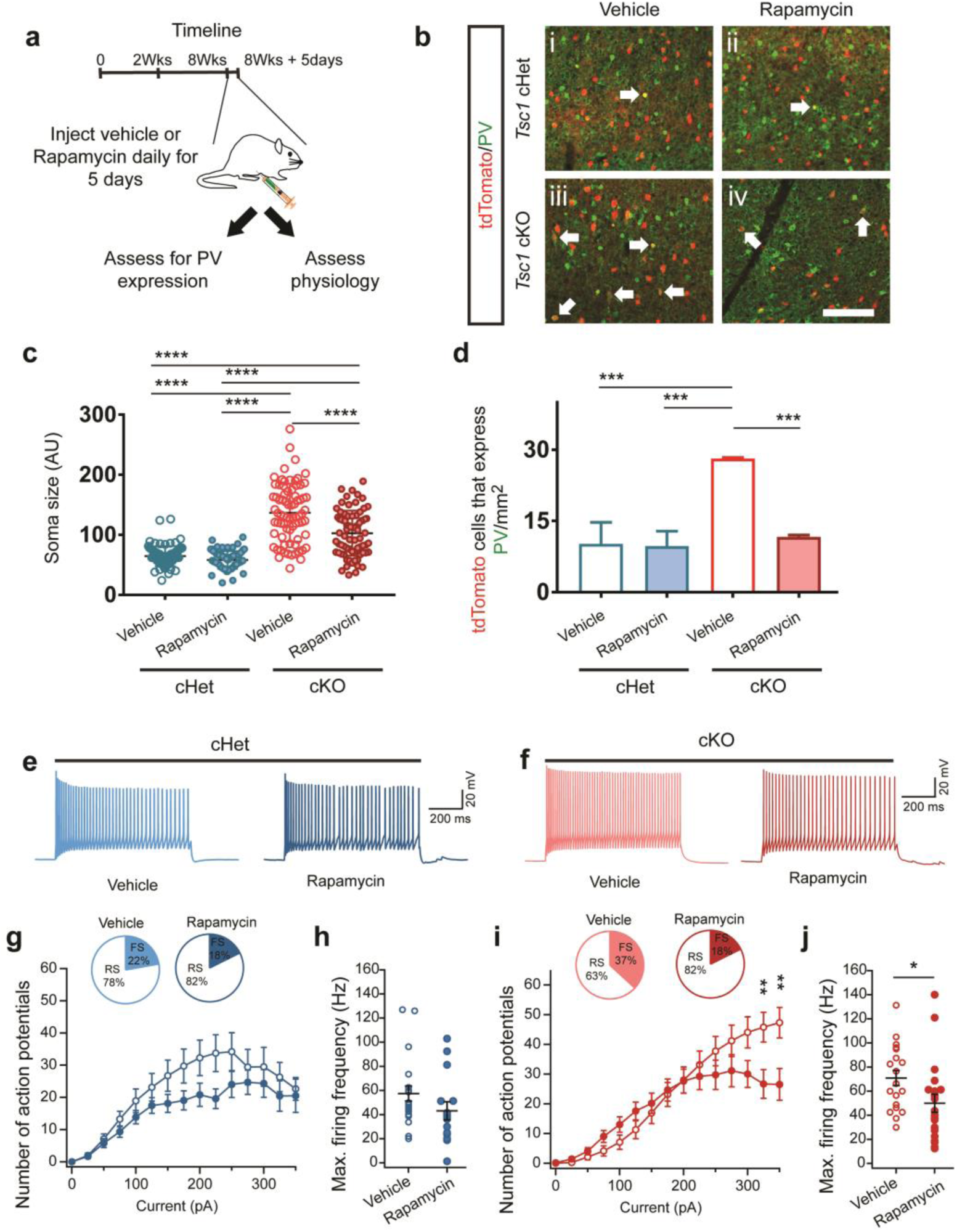
Blocking aberrant MTOR activity with rapamycin decreases PV expression and fast-spiking phenotype in *SST-Cre*-lineage CINs in cKO mice. **(a)** Eight-week old mice that were either *SST-Cre; Tsc1* cHets or cKOs were injected with vehicle or rapamycin consecutively for five days and then assessed a day after for PV expression, cell size and physiological properties. **(b)** Coronal sections were visualized for tdTomato and PV expression from vehicle and rapamycin treated cHets **(i**,**ii)** and cKOs **(iii, iv)**. Arrows point to co-labeled cells. Scale bar is 100µm. **(c)** Quantification of tdTomato+ cell soma size: One-way ANOVA (F_3, 296_ = 95.39, P < 0.0001; cKO vs. all groups, cKO veh vs. rapa, and cHet rapa vs. cKO rapa, p < 0.0001), n = 3 mice per group (25 cells measured from each). (AU) arbitrary units. **(d)** Quantification of the cell density of tdTomato+ cells that co-express PV: One-way ANOVA (F_3, 8_ = 30.51, P = 0.0001; cKO veh vs. cHet veh and cHet rapa, p = 0.0002 and cKO veh vs. cKO, p = 0.0004), n = 3 mice per group. (**e-f**) Example voltage traces in response to suprathreshold current injections in SST+ CINs from vehicle and rapamycin treated cHet **(e)** and cKO **(f)** mice. **(g–h)** Comparison of firing output: Two-way ANOVA: (F_14,475_ = 0.63, P = 0.83); and maximum firing frequency: Two-way unpaired t test (t_33_= 1.4, p = 0.15) of tdTomato+ CINs in vehicle and rapamycin treated cHet mice (Vehicle, n = 18 from 2 mice; Rapamycin, n = 17 cells from 2 mice). Inset, pie charts showing the proportion of SST+ CINs with RS and FS physiological properties in each treatment group. **(i–j)**, Same as **(g–h)** for vehicle and rapamycin treated cKO mice. Note the CINs from rapamycin treated cKO mice have lower firing output: Two-way ANOVA with Bonferroni’s post-test (F_14,555_ = 2.86, P = 0.003; 325 pA, p = 0.004 and 350 pA, p = 0.001), and maximum firing frequency: Two-way unpaired t test (t_37_= 2.1, p = 0.04); (Vehicle, n = 19 cells from 2 mice; Rapamycin, n = 20 from 2 mice). Inset, pie charts showing the proportion of SST-lineage CINs with FS and RS physiological properties in each treatment group. Data are presented as mean ± SD in **(c)** and ± SEM in all other graphs. * p < 0.05, ** p < 0.01, **** p < 0.0001.

As expected, vehicle treated cKO CINs exhibited increased soma size and PV expression compared to the cHet groups (Figs. 7b-d). Rapamycin treatment significantly reduced soma size of SST-lineage CINs in cKOs but not in the cHets (Figs. 7b, c). We also observed reduced PV expression in rapamycin-treated cKOs (Figs. 7b, d). However, rapamycin treatment did not alter PV expression in *SST-Cre*-lineage cHets (Figs. 7b, d).

Next, we tested whether the rapamycin-mediated reduction in PV expression was accompanied by a reduction in FS physiological properties within cKO *SST-Cre*-lineage CINs. Rapamycin treatment did not change the R_in_, firing output or AP properties of cHet CINs (Figs. 7e, g, h, Supplementary Fig. 8). However, in the cKO group, rapamycin treatment did increase R_in_, reduce the rheobase and increase spike frequency accommodation (SFA) (Supplementary Fig. 8). Furthermore, rapamycin reduced the firing output and decreased the fraction of *SST-Cre*-lineage CINs that were classified as FS in cKOs (Figs. 7f, i, j). Notably, even in rapamycin-treated cKO mice, the fraction of FS *SST-Cre*-lineage CINs was higher than the fraction of FS *SST-Cre*-lineage FS CINs in WT mice (Fig. 2h, inset). Rapamycin-treatment also failed to reverse some effects of *Tsc1* loss, including altered AP properties in cKOs (Supplementary Fig. 8). Overall, the five-day rapamycin treatment in adult cKO mice decreased the aberrant PV expression and partially reversed shifts towards FS phenotypes in *SST-Cre*-lineage CINs.

## Discussion

Imbalances in neuronal excitation/inhibition (E/I) are implicated in neuropsychiatric disorders ^11,12,45,46^, and changes in neuronal numbers, synaptic connectivity and intrinsic properties of GABAergic interneurons are, sometimes as a result of compensatory mechanisms, critical contributors to shifts in E/I balance ^13,28,47–50^. TS-Associated Neuropsychiatric Disorders (TANDs) are common manifestations of the disorder ^10^ but the mechanisms underlying TANDs are poorly understood. In this study, we found that deletion of the TS causative gene, *Tsc1*, induced molecular and physiological changes in CINs that may underlie some neuropsychiatric symptoms and seizures observed in TS, (model, supplementary Fig. 9). First, in both *Tsc1* cHet and cKO mice, SST-lineage CINs shifted towards PV-like/FS properties. This reveals a novel mechanism involved in regulating molecular properties and physiology of CINs. Second, loss of *Tsc1* in SST*-*lineages resulted in a neocortex with less synaptic inhibition onto deep layer pyramidal neurons, which may contribute to the higher seizure susceptibility observed in TS. Third, the shift of SST-lineage CINs towards PV expression and FS properties could be reversed, at least partially, by rapamycin administration in adulthood, suggesting that these phenotypes remain plastic in post-mitotic CINs and could provide insights into potential therapeutic mechanisms to treat TS.

Previous studies in a mouse model of TS emphasized that dysregulation of MTOR signaling primarily affects excitatory circuits ^14^, and alters E/I balance through effects on excitatory synapses. While one report examined CINs with regards to TS ^51^, little was known about changes in inhibitory circuits. In this context, we assessed cellular and physiological properties of CINs and inhibitory synapses in *Tsc1* cHets and cKOs. We discovered that loss of *Tsc1* in SST-lineages resulted in aberrant expression of PV and the voltage-gated potassium channel, Kv3.1, in a subset of SST-lineage CINs. This, in turn, shifts their physiological properties from RS to a continuum encompassing both RS and FS properties. Notably, not all SST-lineage CINs adopted these properties, showing that some are less susceptible. This is particularly interesting as recent data has shed new light on subpopulations of SST+ CINs, with distinct molecular identities and functions ^52–54^. Future studies should aim at determining whether the cohort of SST-lineage CINs in which MTOR signaling normally represses PV and Kv3.1 expression as well as FS properties, represents a subtype with specific molecular and/or functional properties. *Tsc1* cHets exhibited similar (but less severe) phenotypes, suggesting that these findings are relevant to TS. Moreover, *TSC1*/2 homozygous mutants have been identified in neural progenitors via a combination of germline and somatic mutations in a TS patient ^55^, suggesting the more severe phenotypes we observed in *Tsc1* cKO mice may also be relevant to some cases of TS.

This study suggests new ideas about how some aspects of CIN cell fate and function are regulated, particularly in post-mitotic/maturing CINs. Previous studies identified TFs that are expressed in SST+ CINs but not in PV+ CINs, as well as TFs that can alter the ratios of SST+ and PV+ CINs ^22–24,27,56–58^. Of note, we are not aware of a TF that is uniquely expressed in PV+ but not SST+ CINs ^59^. One hypothesis to explain this is that PV+ CIN identity is a default state of MGE-derived CINs, and this fate is repressed by TFs restricted to SST-lineages ^59^. Furthermore, perhaps during the maturation of PV-lineage CINs, MTOR activity is induced to promote PV expression and FS properties, i.e. when PV expression first becomes evident during adolescence. This may be achieved as a consequence of activating certain growth factor receptors or potentially via synaptic activity ^7,60^in prospective PV-lineages. However, given that *Tsc1/2* and other components of the MTOR pathway are expressed in most cell types, there would need to be unique factors coupling MTOR signaling in PV-lineages that are different in SST-lineages. These ideas will be investigated in future studies.

Most PV+ CINs have FS physiological properties. In contrast, most SST+ CINs exhibit a RS physiology, although a class of “quasi-fast-spiking” SST+ neurons was recently described ^52^. Moreover, these two neuronal groups are believed to play distinct and dissociable roles in circuit function ^61–65^. Our data suggest new possibilities about these classes. First, CINs exhibit intermediate properties after *Tsc1* loss, which fall along a continuum between the classic SST+ and PV+ distinct groups of CINs. Second, the specification of these classes may be more dynamic in later developmental periods than previously thought. In particular, rapamycin was able to reverse the expression of PV and FS properties in mature SST-lineage cKO CINs, suggesting that some aspects of CIN cell fate phenotypes are malleable by signaling-dependent plasticity mechanisms that remain to be elucidated.

The extent to which changes in molecular (expression of PV vs. SST) and physiological (FS vs.RS) properties correlate with changes in cortical function remains a major question. Differential expression of ion-channels, like delayed-rectifying Kv3 channels, underlie the fast-spiking properties of PV+ CINs ^37,66^. Loss of *Tsc1* leads to an increase in the occurrence of “dual identity” CINs, which express both PV and SST. Furthermore, SST-lineage CINs lacking *Tsc1* have increased expression of Kv3.1 and the properties of these CINs are shifted towards a FS physiology. Interestingly, deletion of FMRP, another monogenic ASD-risk gene, inhibits protein translation downstream of MTOR, of which the Kv3.1 channel is known to be a target in auditory brain stem neurons ^47.^ These converging findings suggest that modulators of the Kv3.1 channel could be potential targets for treating cellular and circuit abnormalities in TS. Future work will be necessary to understand whether *Tsc1* deletion induced shifts in CIN cell fate and physiology underlie behavioral abnormalities in mutant mice and humans diagnosed with TS, which may lead to new therapeutic targets.

In summary, we provided evidence that *Tsc1*-inhibition of MTOR represses PV+/FS properties in a cohort of *SST-Cre*-lineage CINs. Notably, it suggests that regulation of some CIN molecular and physiological properties can be initiated by non-transcriptional events, i.e. MTOR signaling. However, whether transcription is targeted downstream of MTOR signaling is not yet known. Moreover, we identified specific alterations in GABAergic CINs and underlying molecular mediators (e.g. MTOR and Kv3.1) that could plausibly contribute to neurocognitive symptoms of TS. Future studies should assess whether similar abnormalities occur in humans diagnosed with TS (e.g. by immuno-histochemistry in post-mortem tissue) and/or can be reversed by rapamycin analogs (e.g. in neurons derived from human induced pluripotent stem cells), to more directly implicate these findings about CINs in the pathogenesis and/or treatment of TS.

## Methods

### Animals

All mouse strains have been published: *Ai14* Cre-reporter ^68^, *Nkx2.1-Cre* ^33^, *PV-Cre* ^40^, *SST-IRES-Cre* ^31^, *Tsc1*^*floxed* 30^. *Tsc1*^*floxed*^ mice were initially on a mixed C57BL6/J, CD-1 background, then backcrossed to CD-1 for at least four generations before analysis. For timed pregnancies, noon on the day of the vaginal plug was counted as embryonic day 0.5. Both male and female mice were assessed. We did not observe any phenotypic differences between genders and combined all data for each genotype. All animal care and procedures were performed according to the Michigan State University and University of California San Francisco Laboratory Animal Research Center guidelines.

### Cell Counting

To determine cell density (cells/mm^2^), we counted the number of cells in a given section and then divided by the area of that region. To calculate the % of tdTomato^+^ cells that co-labeled with specific markers, we divided the number of co-labeled cells by the total number of tdTomato^+^ cells. For cell transplants, all tdTomato^+^ cells were counted in the neocortex from all sections in a rostral to caudal series. For cell fate counts, only transplants where at least 50 tdTomato^+^ cells could be counted were used for analysis. For soma size quantification, the perimeter of each tdTomato^+^ cell’s soma was traced in Image J, and at least 25 cells were measured and averaged for each mouse.

### Electrophysiology

#### Acute cortical slice preparation

Adult mice of either sex (P45–P60 days) were anesthetized with an intraperitonial injection of euthasol and transcardially perfused with an ice-cold cutting solution containing (in mM) 210 sucrose, 2.5 KCl, 1.25 NaH_2_PO_4_, 25 NaHCO_3_, 0.5 CaCl_2_, 7 MgCl_2_, 7 dextrose (bubbled with 95% O_2_–5% CO_2_, pH ∼7.4). Mice were decapitated, the brains removed and two parallel cuts were made along the coronal plane at the rostral and caudal ends of the brains. Brains were mounted on the flat surface created at the caudal end. Approximately, three coronal slices (250 µm thick) were obtained using Vibrating blade microtome (VT1200S, Leica Microsystems Inc.). Slices were allowed to recover at 34 °C for 30 min followed by 30 min recovery at room temperature in a holding solution containing (in mM) 125 NaCl, 2.5 KCl, 1.25 NaH_2_PO_4_, 25 NaHCO_3_, 2 CaCl_2_, 2 MgCl_2_, 12.5 dextrose, 1.3 ascorbic acid, 3 sodium pyruvate.

#### Whole-cell patch clamp recordings

Somatic whole-cell current clamp and voltage clamp recordings were obtained from submerged slices perfused in heated (32–34**°**C) artificial cerebrospinal fluid (aCSF) containing (in mM); 125 NaCl, 3 KCl, 1.25 NaH_2_ PO_4_, 25 NaHCO_3_, 2 CaCl_2_, 1 MgCl_2_, 12.5 dextrose (bubbled with 95% O_2_/5% CO_2_, pH ∼7.4). Neurons were visualized using DIC optics fitted with a 40x water-immersion objective (BX51WI, Olympus microscope). Pyramidal neurons and tdtomato expressing CINs located in layer 5 were targeted for patching. Patch electrodes (2–4 MΩ) were pulled from borosilicate capillary glass of external diameter 1 mm (Sutter Instruments) using a Flaming/Brown micropipette puller (model P-2000, Sutter Instruments). For current clamp recordings, electrodes were filled with an internal solution containing the following (in mM): 120 K-gluconate, 20 KCl, 10 HEPES, 4 NaCl, 7 K_2_-phosphocreatine, 0.3 Na-GTP, and 4 Mg-ATP (pH ∼7.3 adjusted with KOH). Biocytin (Vector Laboratories) was included (0.1–0.2%) for subsequent histological processing. For voltage clamp recordings, the internal solution contained the following (in mM): 130 Cs-methanesulfonate, 10 CsCl, 10 HEPES, 4 NaCl, 7 phosphocreatine, 0.3 Na-GTP, 4 Mg-ATP, and 2 QX314-Br (pH ∼7.3 adjusted with CsOH).

Electrophysiology data were recorded using Multiclamp 700B amplifier (Molecular Devices). Voltages have not been corrected for measured liquid junction potential (∼8 mV). Upon successful transition to the whole-cell configuration, the neuron was given at least 5 min to stabilize before data were collected. Series resistance and pipette capacitance were appropriately compensated, before each recording. Series resistance was usually 10–20 MΩ, and experiments were terminated if series resistances exceeded 25 MΩ.

#### Electrophysiology protocols and data analysis

All data analyses were performed using custom routines written in IGOR Pro (Wavemetrics). Code is available upon request. Resting membrane potential (RMP) was measured as the membrane voltage measured in current clamp mode immediately after reaching the whole-cell configuration. Input resistance (Rin) was calculated as the slope of the linear fit of the voltage–current plot generated from a family of hyperpolarizing and depolarizing current injections (−50 to +20 pA, steps of 10 pA). Firing output was calculated as the number of action potentials (APs) fired in response to 800 ms long depolarizing current injections (25–500 pA). Firing frequency was calculating as the number of APs fired per second. Rheobase was measured as the minimum current injection that elicited spiking. Firing traces in response to 50 pA current above the rheobase were used for analysis of single AP properties– AP threshold, maximum *dV*/*dt* (rate of rise of AP), AP amplitude, AP half-width and fast after hyperpolarization (fAHP) amplitude. Threshold was defined as the voltage at which the value of third derivative of voltage with time is maximum. Action potential amplitude was measured from threshold to peak, with the half-width measured at half this distance. Fast after hyperpolarization was measured from the threshold to the negative voltage peak after the AP. Index of spike-frequency accommodation (SFA) was calculated as the ratio of the last inter-spike interval to the first inter-spike interval. Coefficient of variance (CV) for inter-spike interval (ISI), AP amplitude and AP half-width was calculated as the ratio of standard deviation to the mean. Recorded CINs in all genotypes were classified as fast-spiking or regular-spiking based on electrophysiological properties. Specifically, CINs were classified as fast-spiking if the AP half-width was < 0.5 ms, firing frequency > 50 Hz, fAHP amplitude was > 14 mV and SFA was < 2. The correspondence between PV expression and fast-spiking electrophysiological properties was confirmed in a subset of CINs by quantification of PV staining in biocytin-filled neurons (16 of 21 biocytin-labelled CINs classified as fast-spiking were PV+).

Spontaneous inhibitory currents were recorded for 5 min with neurons voltage clamped at +10 mV. Miniature inhibitory currents were recorded in the presence of TTX. Spontaneous and miniature inhibitory currents were analyzed off-line using Clampfit (pClamp, Molecular devices) event detection.

For Cre-dependent expression, 600 nl of *AAV-DIO-ChR2-eFYP* virus was bilaterally injected into P30–40 *SST-Cre* mice. We waited at least 4 weeks after virus injection before preparing brain slices. To measure optogenetically evoked inhibitory currents (oIPSCs), we voltage-clamped Layer 5 pyramidal neurons at +10 mV. We stimulated ChR2 using 5 ms long single or multiple light pulses (maximum light power, 4 mW/mm^2^) generated by a Lambda DG-4 high-speed optical switch with a 300 W Xenon lamp (Sutter Instruments) and an excitation filter centered around 470 nm, delivered to the slice through a 40X objective (Olympus). Frequency dependent attenuation (20 Hz) and paired-pulse ratios of oIPSCs were measured at the maximum light power.

To measure unitary inhibitory currents (uIPSCs), dual whole cell recordings were obtained from connected layer 5 tdTomato expressing (*SST-Cre* lineage or *PV-Cre* lineage) CINs and adjacent layer 5 pyramidal neurons. We obtained current clamp recordings from tdTomato+ CINs and voltage clamp recordings (+10 mV) from pyramidal neurons. Short duration (1–2 ms) depolarizing current steps (1–1.5 nA) were injected into the CINs to elicit single action potentials and uIPSCs were recorded in postsynaptic pyramidal neurons. Average of 20–30 uIPSCs were used to calculate the peak amplitude (from the baseline to the peak of uIPSC), rising slope (slope of a fitting line from 10–90% after onset of uIPSC) and total charge of uIPSCs (integral of uIPSCs from the onset until the trace returned completely to the baseline).

### Computer simulations

Simulations were performed using the NEURON 7.4 simulation environment (Carnevale and Hines, 2006) with Python interface. Four different morphologically detailed models of SST+ neocortical layer 5 CINs were used from Allen Brain Atlas; Allen Institute (2015) Documentation, http://help.brain-map.org/display/celltypes/Documentation. Model files for simulation in Python environment were downloaded from ModelDB (senselab.med.yale.edu). Maximum conductance densities (in S/cm^2^) of Kv3.1 channel (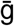_kv3.1) were noted for the four different models (0.25, 0.009, 0.29, 0.56). These values were used for the low Kv3.1 conductance conditions. To simulate the effects of increase in Kv3.1 expression, 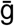_kv3.1 was increased to 1.5 in all four models. Firing output in the low and high Kv3.1 conditions were measured as the number of spikes fired in response to 1 s long depolarizing current steps (50–350 pA). Spike times were measured by determining the time of AP peak. Instantaneous frequency (1/inter-spike interval) was computed from current injections that elicited >40 APs. Subthreshold membrane properties were calculated from the voltage traces in response to hyperpolarizing and depolarizing current injections (−50 to +20 pA, steps of 10 pA).

### Immuno-fluorescent tissue staining

Immunofluorescent labeling was performed on 25µm cryo-sectioned tissue with the following antibodies: mouse anti-Kv3.1b 1:400 (NeuroMab cat. # 75-041); rabbit anti-PV 1:400 (Swant cat. # PV 27); rat anti-SST 1:200 (Millipore cat. # MAB354). The appropriate 488, 594 or 647 Alexa-conjugated secondary antibodies were from Thermo Fisher Scientific (1:300 dilution). Sections were cover-slipped with Vectashield (Vector labs).

### MGE cell transplantation

Lentiviral transduction of MGE cells followed by intracranial transplantation was performed as previously described ^43^. First, HEK293T cells were transfected using Lipofectamine2000 (Thermo Fischer Scientific) with four plasmids to generate lentivirus particles. E13.5 MGEs from individual embryos were dissected in ice-cold HBSS and then kept on ice in DMEM media (containing 10% fetal bovine serum). MGEs were then mechanically dissociated with a p1000 pipette tip and transduced with lentivirus. To perform transductions, dissociated MGE cells were mixed with pre-warmed media that was at physiological pH and was incubated with ∼10-20 μls of concentrated lentivirus, with titers ranging from 1X10^8-1X10^10 infectious units/ml. They were next incubated at 37°C for 30 minutes, with intervals of agitation. Since the MGE cells had a Cre-dependent reporter, *Ai14* ^68^, only MGE cells transduced with Cre-expressing lentiviruses were visible in the host neocortex after transplantation. This combinatorial method allows the transduced/transplanted cells to express a strong reporter, i.e. tdTomato expressed from the beta-actin promoter, while maintaining cell type specificity. Cells were then pelleted in a tabletop centrifuge at low speed (700xg, ∼4 minutes) and washed 3 times with media followed by trituration to disperse cells between each wash to remove excess virus. The final pellet was left under a few μls of media, put on ice, and the remaining media was removed with a fine point kim wipe before the injection needle was loaded. For injections, a glass micropipette with a 45° beveled tip, ∼50 μm outer diameter, was preloaded with sterile mineral oil and cells were then front-loaded into the tip of the needle using a plunger connected to a hydraulic drive (Narishige MO-10) that was mounted to a stereotaxic frame. Pups were anesthetized on ice for 1-2 minutes before being placed on a mold for injections. Each pup received 3-6 injections of cells, at 70 nl per site, in their right hemisphere. These sites were about 1 mm apart from rostral to caudal and medial to lateral and were then injected into layers V-VI of a P1 WT neocortex. After injections, pups were allowed to recover and then put back with their mother. The transplanted mice were sacrificed at 35 days post-transplant and transcardially perfused with PBS followed by 4% PFA. Brains were then post-fixed in 4% PFA for a short interval, ∼30 minutes, and then sunk in 30% sucrose before embedding in OCT.

#### DNA vector generation

The *DlxI12b-BG-IRES-Cre* lentiviral vector was previously described (Vogt et al., 2017). To generate the *DlxI12b-BG-hTSC1-IRES-Cre* lentiviral vector, the human *TSC1* gene was PCR amplified from a vector containing the *hTSC1* cDNA (Addgene) ^70^. The primers (Forward 5’-GAGA<u>GCTAGC</u>ATGGCCCAACAAGCAAATG-3’ and Reverse 5’-GAGA<u>TGTACA</u>TTAGCTGTGTTCATGATG-3’), were used to introduce NheI and BsrGI restriction enzyme sites, respectively (underlined). Next the PCR product and the *DlxI12b-BG-IRES-Cre* vector were digested with the appropriate enzymes and ligated. The vector was verified by restriction digest and sequencing.

### Rapamycin treatment

Rapamycin (LC Laboratories, Woburn, MA, USA) was dissolved in ethanol at a concentration of 20 mg/ml and stored at −20°C. On the day of the injections, drug was resuspended in saline (containing 0.25% PEG and 0.25% Tween-80). Mice received rapamycin solution (10 mg kg^-1^) or an equal volume of vehicle by intraperitoneal injection once daily for five consecutive days.

### Quantification and statistical analysis

For anatomical analyses, statistics were performed using Prism version 6, a p value of < 0.05 was considered significant. For most parametric measurements, we used a One-Way ANOVA with Tukey’s post test to determine significance (cell density and soma size measurements). For non-parametric data sets, we used a Chi-squared test with Yate’s correction to determine significance (data normalized and expressed as a percentage). For electrophysiological analyses, statistical comparisons were done using custom-written software in MATLAB or Prism (Graphpad). Statistical significance was assessed using a paired or unpaired two-tailed Student’s t-test. For repeated observations, two-way ANOVA followed by a post hoc Tukey’s or Bonferroni’s test was used. Statistical comparisons across three groups were done using one-way ANOVA followed by a post hoc Tukey test. Probability values for ANOVAs are denoted as “P” and the probability values for post hoc comparisons are denoted as “p”. Error bars indicate ± SEM unless otherwise noted.

## Acknowledgements

**VSS** was funded by CDMRP (#TS150059) and NIMH R01 (#MH106507). **JLR** by Nina Ireland, NIMH R01 (#MH081880), NIMH R37 (#MH049428) and CDMRP (#TS150059). **RM** by NARSAD Young Investigator Award (Leichtung Family Investigator). **ELLP** by UCSF Neuroscience Program and NIMH R01 (#MH081880). **ANR** by NIMH R01 (#MH081880). **AMS, KA** and **DV** by the Alliance Funds from Michigan State University/Helen DeVos Children’s Hospital.

