## Supplemental material for "The Tuberous Sclerosis gene, *Tsc1*, represses parvalbumin+/fast-spiking properties in somatostatin-lineage cortical interneurons"

### Supplemental materials and methods

#### Fluorescent *in situ* hybridization

Brain tissues from P30*SST-CRE* WT and *Tsc1* cKOs were perfused and incubated with 4% PFA for 1 hour, followed by incubation in 30% sucrose/PBS for cryoprotection until the day of section. Tissues were embedded in OCT and cryo-sectioned to generate 40um tissue sections.

To generate the *Tsc1* DNA vector and riboprobe, *Tsc1* cDNA was PCR amplified from homemade mouse cDNA library synthesized from P0 CD-1 mouse neocortex using Superscript II.

The following primers were used:

5' GAGA<u>ATCGAT</u>AGACAAAGCTGGAGGACTGC

3' ATAT<u>TCTAGA</u>TCAAGCCTCTCTTCTGCTGC

ClaI and XbaI restriction enzymes sites were introduced (underlined). The primers target the *Tsc1* floxed regions, which flank exons 16 and 17. Next, the *Tsc1* PCR product and the vector, pSP73 (Promega Cat # P2221), were digested with ClaI and XbaI, and then ligated. The *Tsc1* RNA anti-sense fluorescein-labeled probe was generated by T7 RNA polymerase (Roche) and the Fluorescein labeling kit (Roche) from a NdeI linearized vector, with the size of the probe ~600bp. *Sst* Digoxigenin-labeled riboprobe was kindly provided by Dr. John Rubenstein (original resource T. Lufkin). Fluorescent in situ hybridization was performed as previously described [1]. Imaging was done with a 60x objective (Nikon Apo 1.4 oil) using a Nikon Ti microscope with DS-Ri2 color camera.

#### Morphological reconstructions

In all our recordings, 0.2–0.3% biocytin was added to the pipette solution. Molecular identity and morphological characteristics of recorded cells were confirmed by tdTomato expression and subsequent staining for PV and biocytin. Slices containing biocytin-filled cells were fixed overnight in a buffered solution containing 4% paraformaldehyde. Slices were rinsed twice in PBS, then blocked and permeabilized for 3hr in PBS with 10% FBS, 0.5% Triton X-100 and 0.05% sodium Azide. Slices were immuno-stained overnight with primary antibody: rabbit anti-PV (Swant, cat. # PV27) diluted 1:1000 in PBS with 0.1% Triton X-100, 10% FBS and 0.025% sodium azide. Slices were washed 2 x 30min in PBS with 0.25% Triton X-100, and 2 x 30min in PBS. Goat anti-rabbit Alexa-488 secondary antibody (1:750, Thermo Fisher, cat. #A-11034) and Streptavidin-647 (1:500, Thermo Fisher, cat. # S-32357) were added with Hoescht 33342 nuclear counterstain (1:2000, Thermo Fisher, cat. # H3570) for 4-6hr at room temperature, then overnight at 4°C. After washing 2 x 30min in PBS with 0.25% Triton

X-100, and 2 x 30min in PBS, slices were mounted on Superfrost Plus slides and coverslipped with Dako Fluorescence Mounting Medium (cat. # S3023).

Low-magnification epifluorescent images were taken using a Coolsnap camera (Photometrics) mounted on a Nikon Eclipse 80i microscope using NIS Elements acquisition software (Nikon). Confocal images were taken with 20x air and 40x oil objectives on an Andor Borealis CSU-W1 spinning disk confocal mounted on a Nikon Ti Microscope (UCSF Nikon Imaging Center, NIH S10 Shared Instrumentation grant 1S10OD017993-01A1) and captured with an Andor Zyla sCMOS camera and Micro-Manager software (Open Imaging). Confocal stacks were imported in NeuTube for semi-automated tracing of biocytin-filled neurons [2]. Reconstructions were analyzed using Trees toolbox in MATLAB [3].

Phosphorylated ribosomal subunit S6 immunofluorescent labeling

P35 frozen brain sections, 25µm thickness, were stained with a rabbit anti-phosphoS6$^{SER240/244}$, (1:400 Cell Signaling Technologies, cat. # 5364). Briefly, sections were washed 3x in PBS containing 0.3% Triton X-100 (PBST) and then blocked for 1 hour in the same solution supplemented with 5% bovine serum albumin. The antibody was applied for 2 hours at room temperature (RT), then washed 3x with PBST. Secondary donkey anti-rabbit Alexa 488 (1:300, Thermo Fisher, cat. # A-21206) was applied for 1 hour at RT then the sections were washed 3x in PBST before slides were mounted with Vectashield (Vector labs).

### Supplemental Figures

#### Supplementary Fig. 1

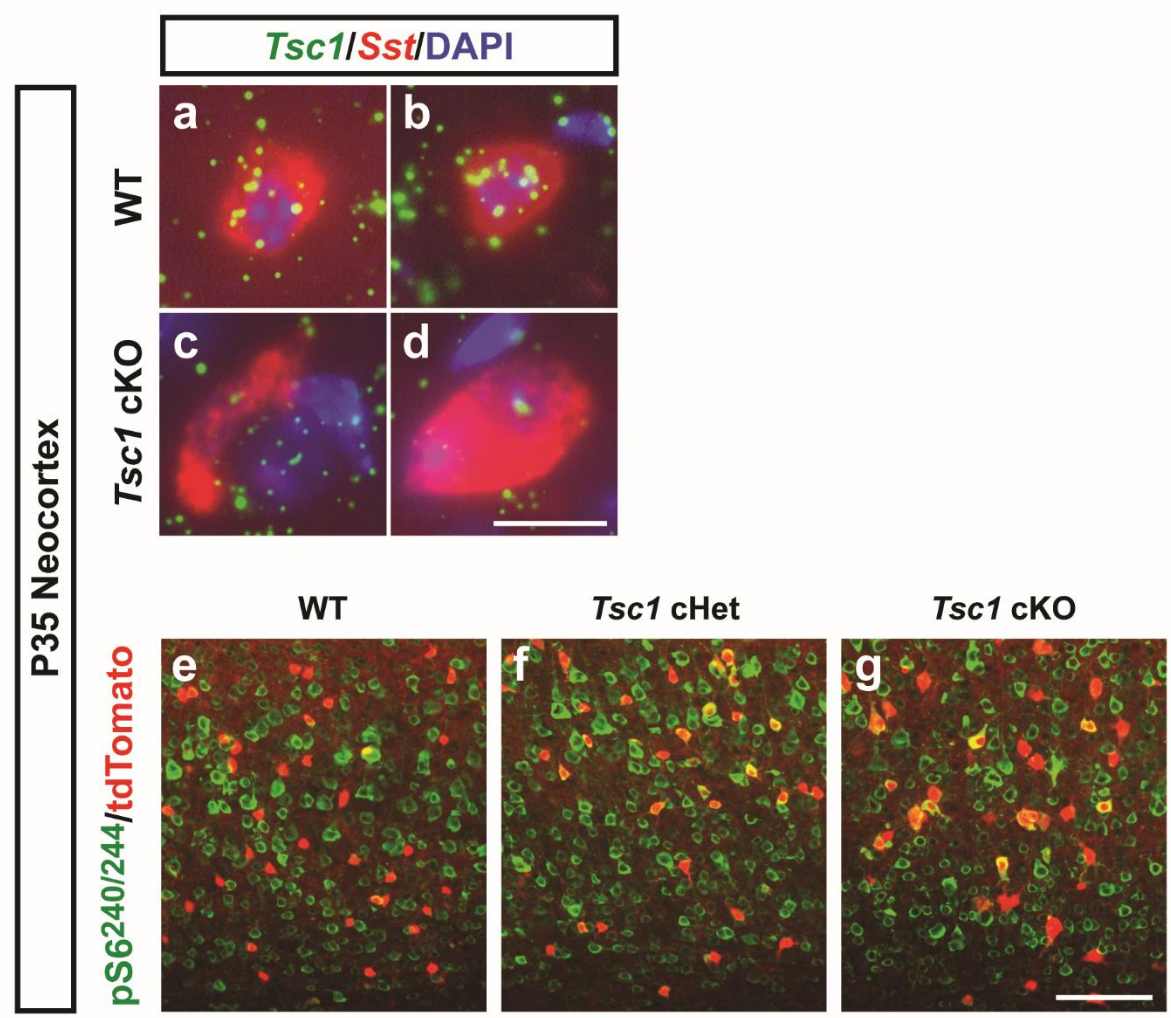


**Validation of *Tsc1* loss in *SST-Cre*-lineage CINs. (a-d)** *Tsc1* (green) and *Sst* (red) fluorescent *in situ* hybridization from WT **(a-b)** and *Tsc1* cKO **(c-d)** P35 neocortex. Note the decreased expression of *Tsc1* and increased cell soma size in the *Tsc1* cKO. **(e-g)** *SST*-*Cre*-lineage CINs (tdTomato+) co-labeled for the MTOR target, phosphorylated ribosomal subunit S6 at serines 240 and 244 (green). Scale bars in **(d)** = 20µm and **(g)** = 100µm.

#### Supplementary Fig. 2

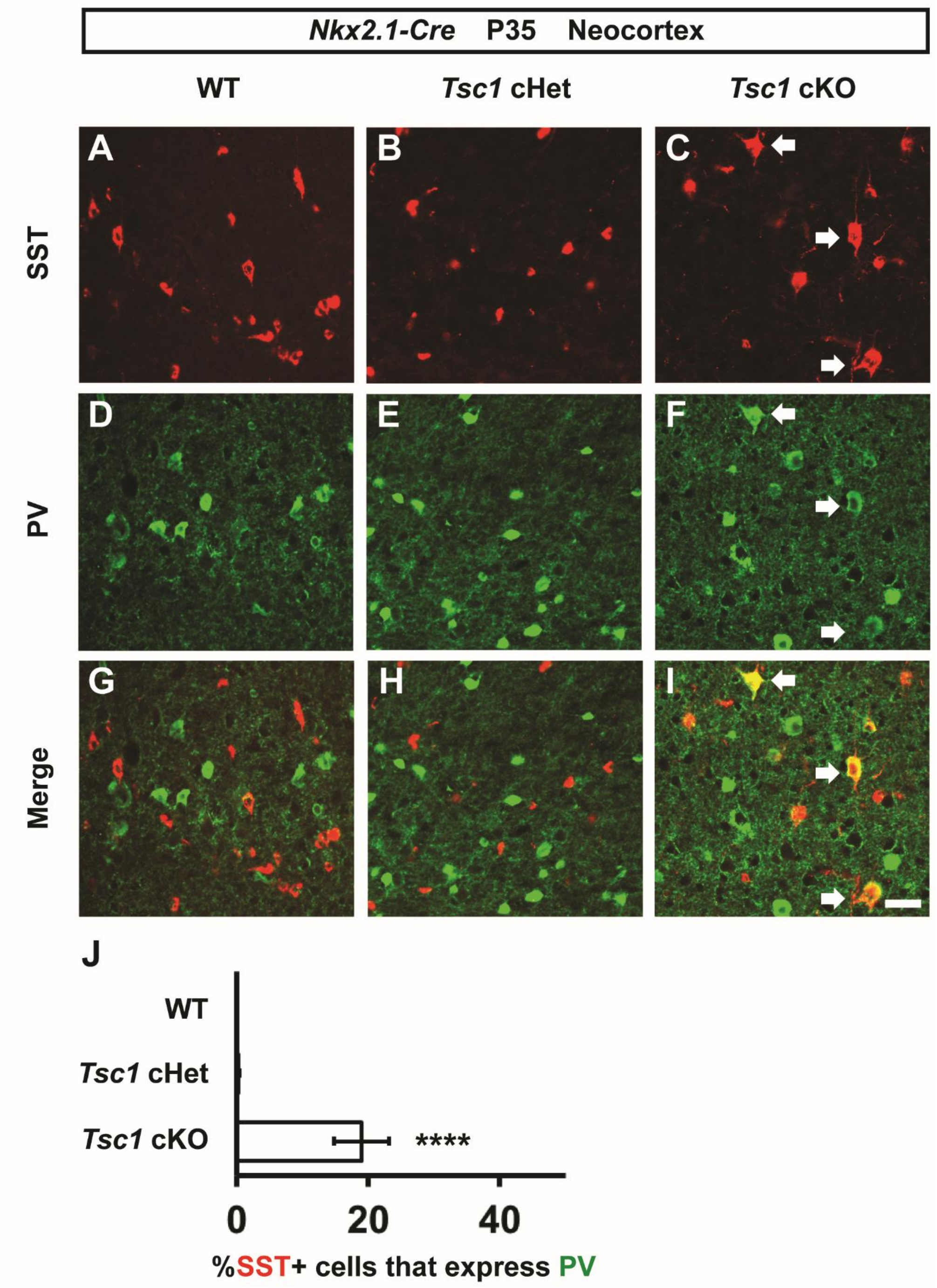


**Co-expression of SST and PV in the somatosensory cortex of *Nkx2.1-Cre; Tsc1* cKOs.** P35 coronal immunofluorescent tissue sections were assessed for somatostatin (SST) **(a-c)** and parvalbumin (PV) **(d-f)** in the somatosensory cortex. Merged images did not reveal any co-expression in WT **(g)** and few in *Tsc1* cHet **(h)** tissue, however, *Tsc1* cKO tissue had multiple co-expressing cells **(i)**. Arrows point to co-expressing cells. (j) Quantification of the % of SST+ CINs co-labeled for PV: (Chi-squared test with Yate's correction **** $p < 0.0001$, n=3, all groups). Data are expressed as the mean ± SEM. Scale bar in **(i)** = 100 µm. **** $p < 0.0001$.

#### Supplementary Fig. 3

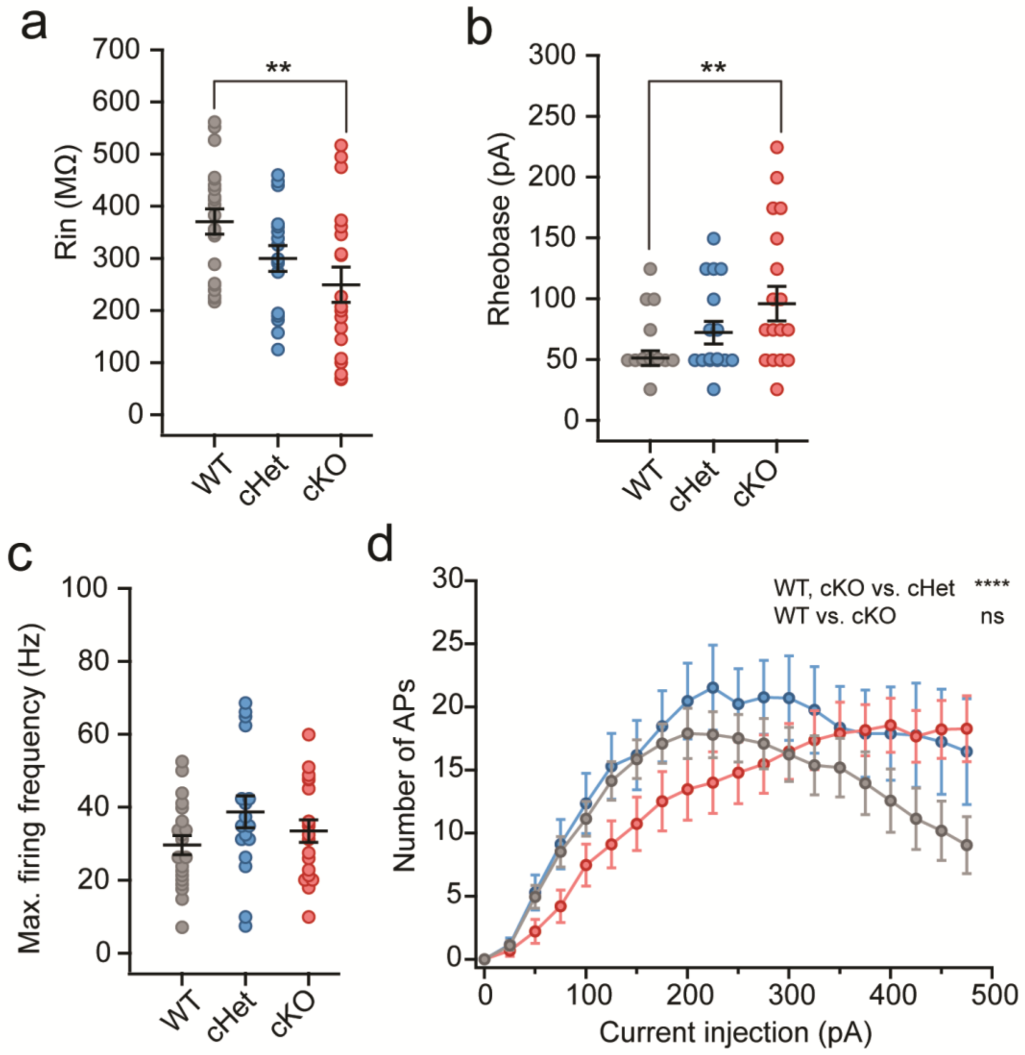


**Increased proportion of *SST-Cre*-lineage CINs with fast-spiking firing properties underlies the increased firing output in cKO group.** Comparison of input resistance ($R_{in}$): One-way ANOVA ($F_{2, 54} = 5.02$, P = 0.007) **(a)**; rheobase, One-way ANOVA ($F_{2, 54} = 5$, p = 0.007, WT vs cKO, p = 0.007)) **(b);** and maximum firing frequency, One-way ANOVA ($F_{2, 54} = 1.77$, WT vs cKO, p = 0.007) **(c)** of regular-spiking SST-lineage CINs in WT (grey), cHet (blue) and cKO (red) groups. Note the $R_{in}$ of regular-spiking CINs in cKO group is significantly lower and rheobase is higher as compared to WT group (WT, n=21 cells from 4 mice; cHet, n=17 cells from 3 mice; cKO, n=19 cells from 4 mice). **(d)** Number of APs fired in response to steps of depolarizing current injections: Two way ANOVA ($F_{(2,1079)} = 11.1$, P < 0.0001; WT and cKO vs. cHet, p < 0.0001; WT vs. cKO, p = 0.97). Data are presented as mean ± S.E.M. **p < 0.01, **** p < 0.0001.

#### Supplementary Fig. 4

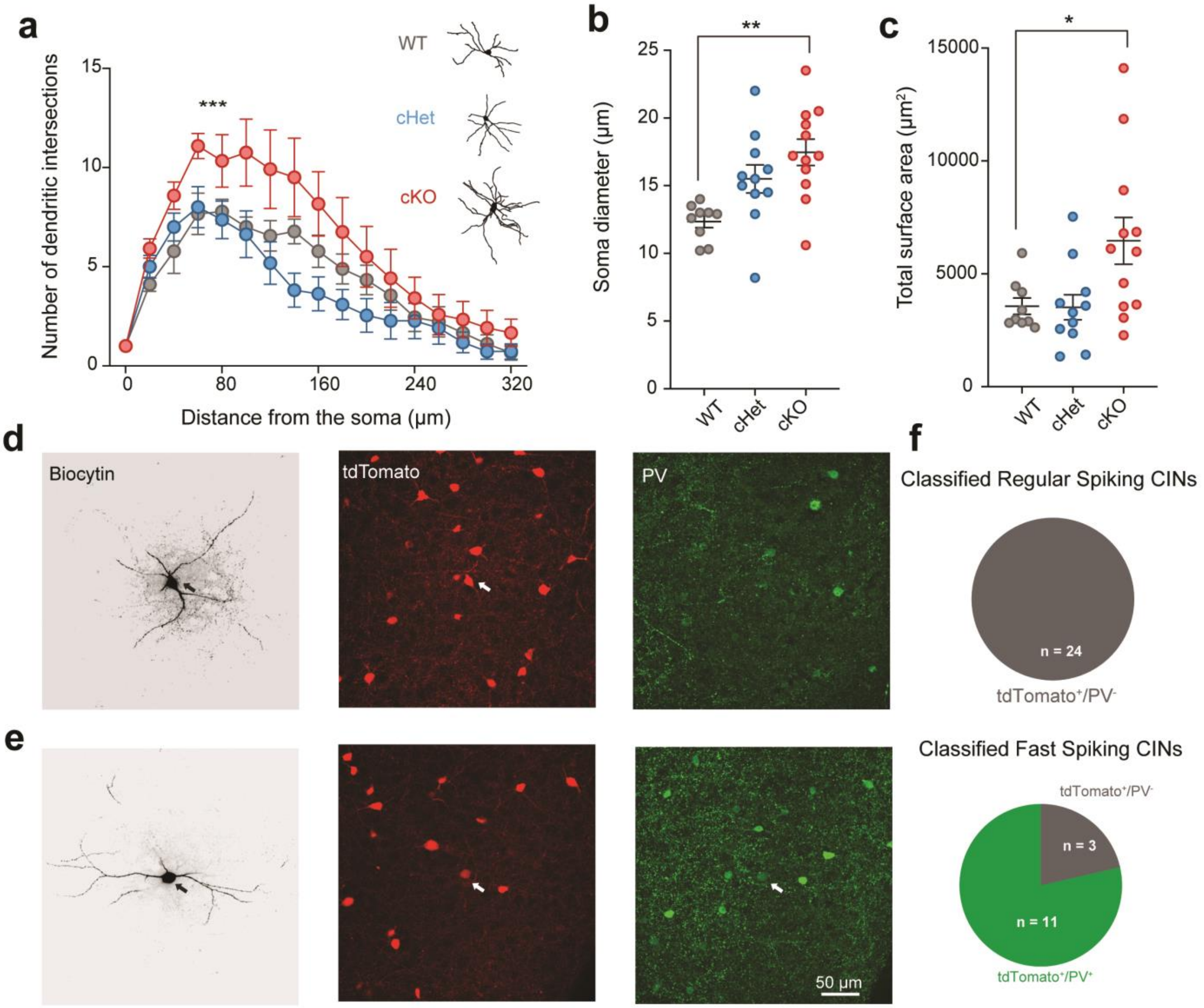


***Tsc1* deletion increases PV expression and dendritic complexity of *SST-lineage* CINs. (a)** Number of dendritic intersections during Sholl analysis. Note, the cKO CINs had significantly more intersections (Repeated measures ANOVA, Interaction, $F_{32, 464} = 2.15$, P = 0.0004; WT vs. cKO, P = 0.0003; cHet vs cKO, P < 0.0001). **(b)** Soma diameter, One-way ANOVA ($F_{2, 29} = 7.5$, P = 0.002 and WT vs. cKO, P = 0.001). **(c)** Total surface area, One-way ANOVA ($F_{2, 29} = 4.9$, P = 0.013; WT vs. cKO, P = 0.03). WT, n = 9; cHet, n = 11; cKO, n=12**. (d)** Layer 5 *SST-lineage* CINs in *Tsc1* cKOs were filled with biocytin (grey) and later confirmed to express tdTomato (red) but not PV (green). **(e)** Layer 5 *SST-lineage* CIN in cKO filled with biocytin and later confirmed to express both tdTomato and PV. **(f)** Top: Pie chart showing the proportion of biocytin-filled CINs in cKOs and cHets classified as RS that also expressed PV. Bottom: Pie chart showing the proportion of biocytin-filled CINs in cKOs and cHets classified as FS that also expressed PV. Data are presented as mean ± S.E.M. * p < 0.05, **p < 0.01, *** p < 0.001

#### Supplementary Fig. 5

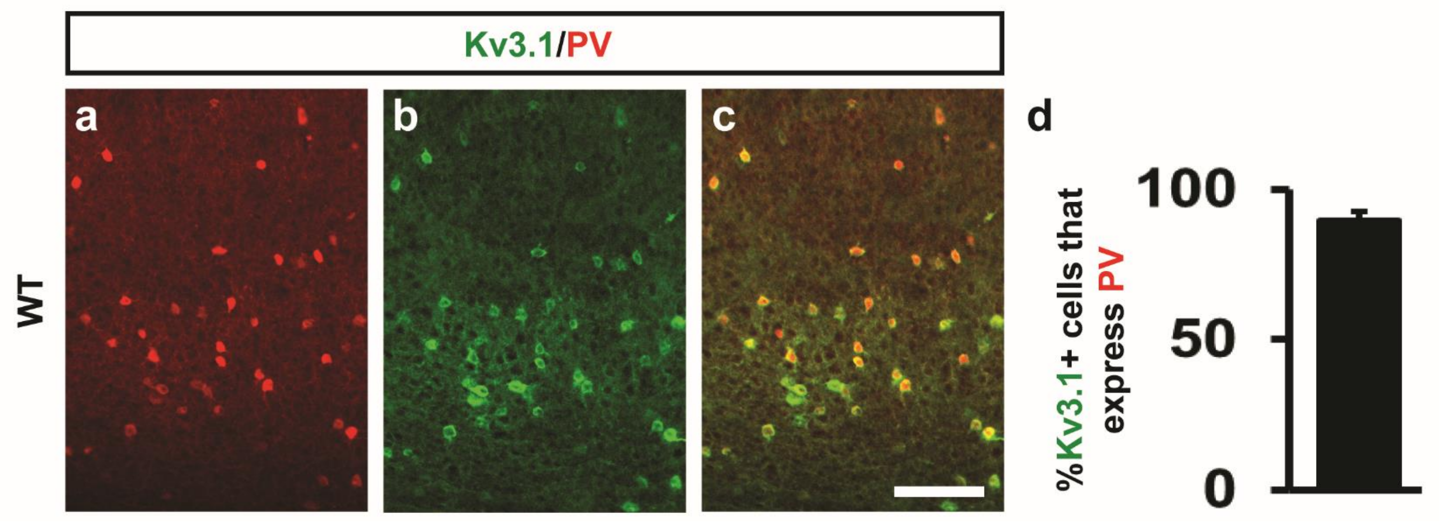


**Validation of preferential expression of Kv3.1 in PV+ CINs. (a-c)** P35 WT neocortices were co-labeled for parvalbumin (PV) and the potassium channel, Kv3.1. Scale bar in **(c)** = 100µm. **(d)** quantification of the proportion of Kv3.1+ cells that express PV. Data are expressed as the mean ± SEM. n = 3.

#### Supplementary Fig. 6

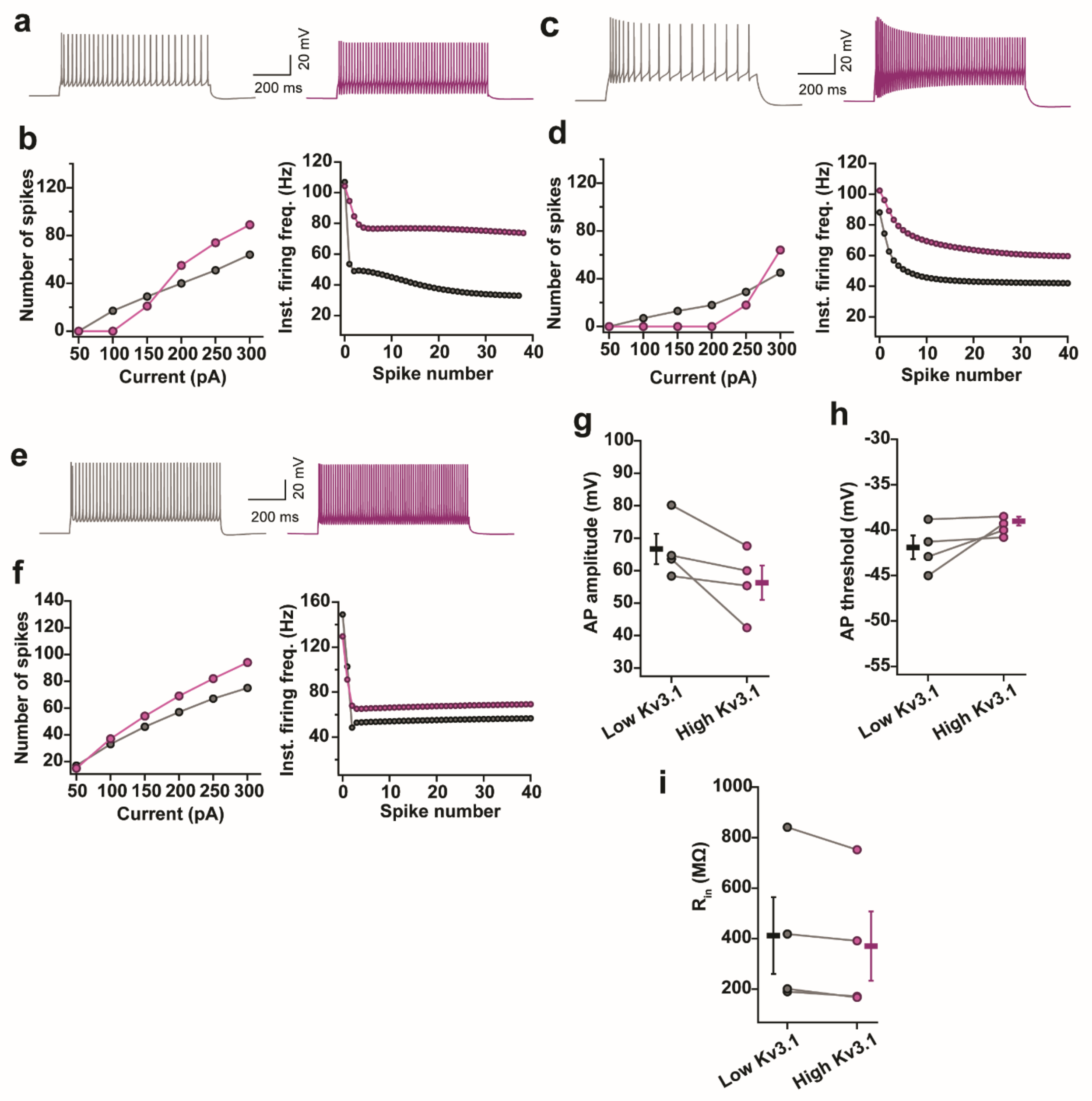


**Increased Kv3.1 conductance switches the physiological properties of SST+ CINs from regular spiking to fast spiking. (a)** Example firing traces from a morphologically realistic model of SST+ CINs with low expression of Kv3.1 (grey) and high expression of Kv3.1 (purple). **(b)** Firing data from one model is shown. Increasing the conductance of Kv3.1 channel increases the number of APs fired in response to depolarizing current injections (left) and decreases the change in instantaneous firing frequency (right) during a train of APs. **(c–f)** Same as **(a, b)** for two other models of SST+ CINs models. **(g–i)** Increasing the Kv3.1 conductance in four different models of SST+ CINs does not affect the AP amplitude (Paired two-tailed t-test, $t_3 = 2.48$), AP threshold (Paired two-tailed t-test, $t_3 = 1.8$) and input resistance ($R_{in}$) (Paired two-tailed t-test, $t_3 = 2.66$). Data are presented as mean ± S.E.M.

#### Supplementary Fig. 7

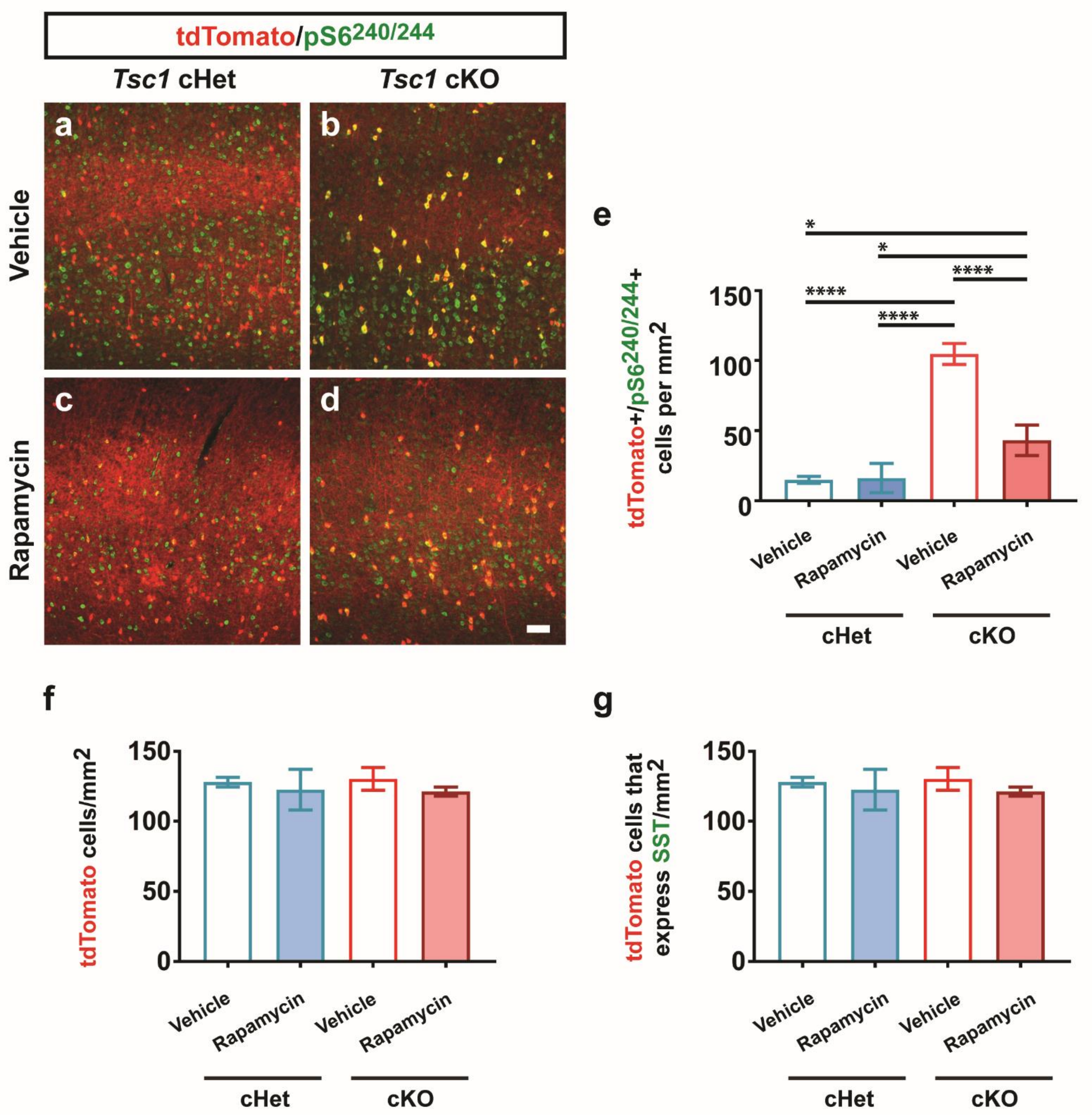


**Validation of rapamycin treated mice and additional cell counts.** Vehicle treated **(a, b)** or rapamycin treated **(c, d)** *Tsc1; SST-Cre* cHet and cKO neocortices were immuno-fluorescently labeled for the MTOR target, phosphorylated ribosomal subunit $S6^{Ser240/244}$. **(e)** quantification of the cell density of *SST-Cre*-lineage cells (tdTomato+) that co-label for pS6 in the neocortex: One-way ANOVA ($F_{3,\,8}$ = 73.3, p < 0.0001; cKO veh vs. all groups P < 0.0001; cKO rapa vs. cHet veh P = 0.02 or cHet rapa P = 0.02). **(f)** quantification of the cell density of tdTomato+ cells: One-way ANOVA ($F_{3,\,8}$ = 0.9). **(g)** quantification of the cell density of tdTomato+ cells that co-express SST: One-way ANOVA ($F_{3,\,8}$ = 1.8). Data are represented as mean ± SEM. Scale bar in **(d)** = 100µm. n = 3, all groups. * p < 0.05, **** p < 0.0001.

**Supplementary Fig. 8**

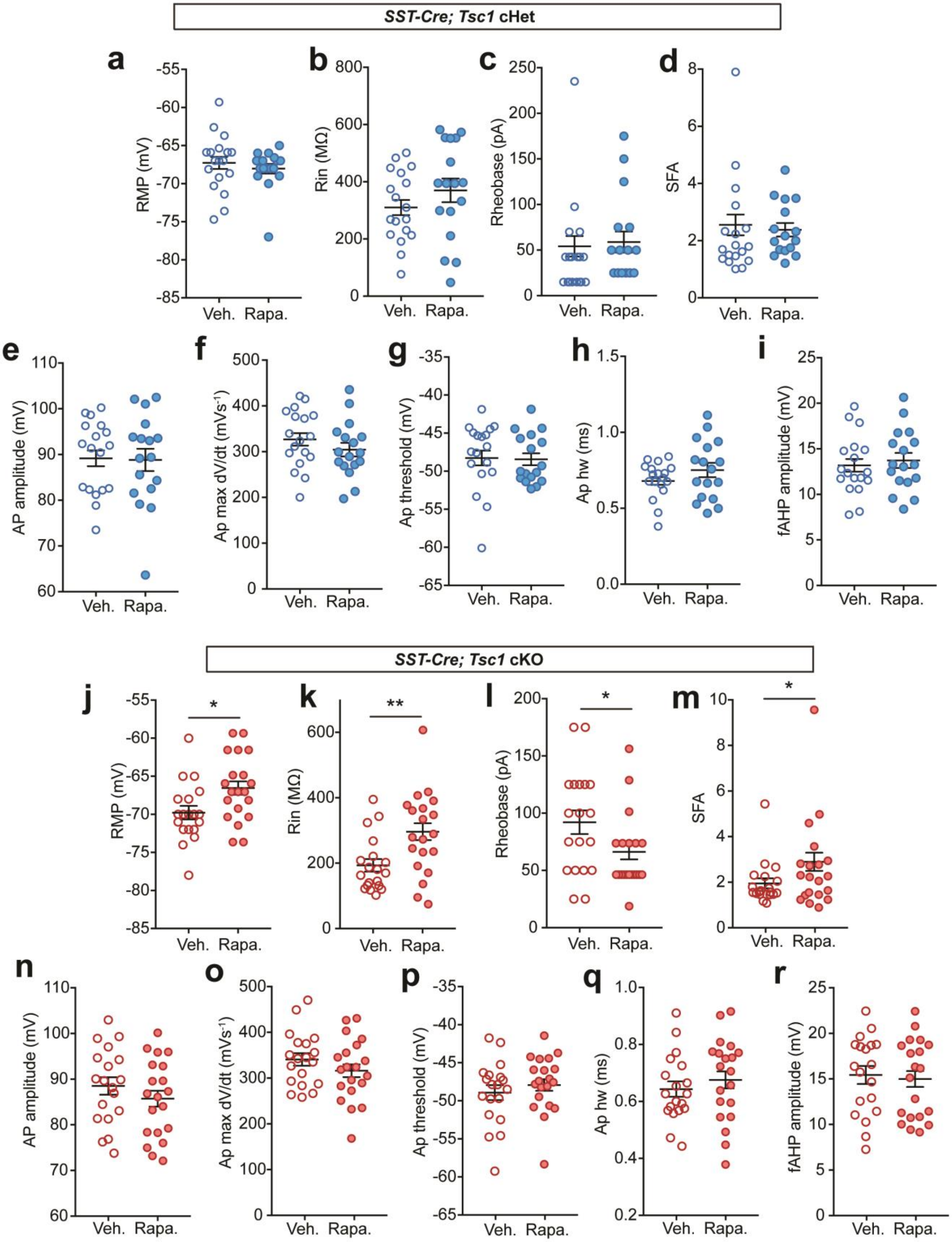

**Blocking aberrant mTOR activity with rapamycin affects the subthreshold and firing properties of SST-lineage CINs in *Tsc1* cKO mice. (a, b)** Resting membrane potential **(**RMP) and input resistance ($R_{in}$) of SST-lineage CINs in vehicle (n = 18 from 2 mice) and rapamycin (n = 17 cells from 2 mice) treated cHets were compared: Unpaired two-way t test; RMP: ($t_{33}$ = 0.73; $R_{in}$: $t_{33}$ = 1.2). **(c, d)** Rheobase and spike-frequency accommodation (SFA) of SST-lineage CINs in vehicle and rapamycin treated cHets were compared: Unpaired two-way t test; Rheobase: ($t_{33}$ = 0.28; SFA: $t_{33}$ = 0.44). **(e–i)** Action potential (AP) amplitude, maximum rate of rise (max dV/dt), threshold, half-width and fast afterhyperpolarization (fAHP) amplitude of SST-lineage CINs in vehicle and rapamycin treated cHets were compared: Unpaired two-way t test; AP amplitude: ($t_{33}$ = 0.11; max dV/dt: $t_{33}$ = 0.12); AP threshold: ($t_{33}$ = 0.13); AP half-width: ($t_{33}$ = 1.3); fAHP: ($t_{33}$ = 0.49). **(j, k)** Resting membrane potential **(**RMP) and input resistance ($R_{in}$) of SST-lineage CINs in vehicle (n = 19 cells from 2 mice) and rapamycin (n = 20 cells from 2 mice) treated cKOs were compared: Unpaired two-way t test; RMP: ($t_{37}$ = 2.5, P = 0.01); ($R_{in}$: $t_{37}$ = 3.1, P = 0.003). **(l, m)** Rheobase and spike-frequency accommodation (SFA) of SST-lineage CINs in vehicle and rapamycin treated cHets were compared: Unpaired two-way t test; Rheobase: ($t_{37}$ = 2.1, P = 0.039); SFA: ($t_{37}$ = 1, p = 0.04). **(n–r)** Action potential (AP) amplitude, maximum rate of rise (max dV/dt), threshold, half-width and fast afterhyperpolarization (fAHP) amplitude of SST-lineage CINs in vehicle and rapamycin treated cHets were compared: Unpaired two-way t test; AP amplitude: ($t_{37}$ = 1); max dV/dt: ($t_{37}$ = 1.2); AP threshold: ($t_{37}$ = 0.8); AP half-width: ($t_{37}$ = 0.8); fAHP: ($t_{37}$ = 0.33). Data are presented as mean ± S.E.M; * $p < 0.05$, ** $p < 0.01$.

#### Supplementary Fig. 9

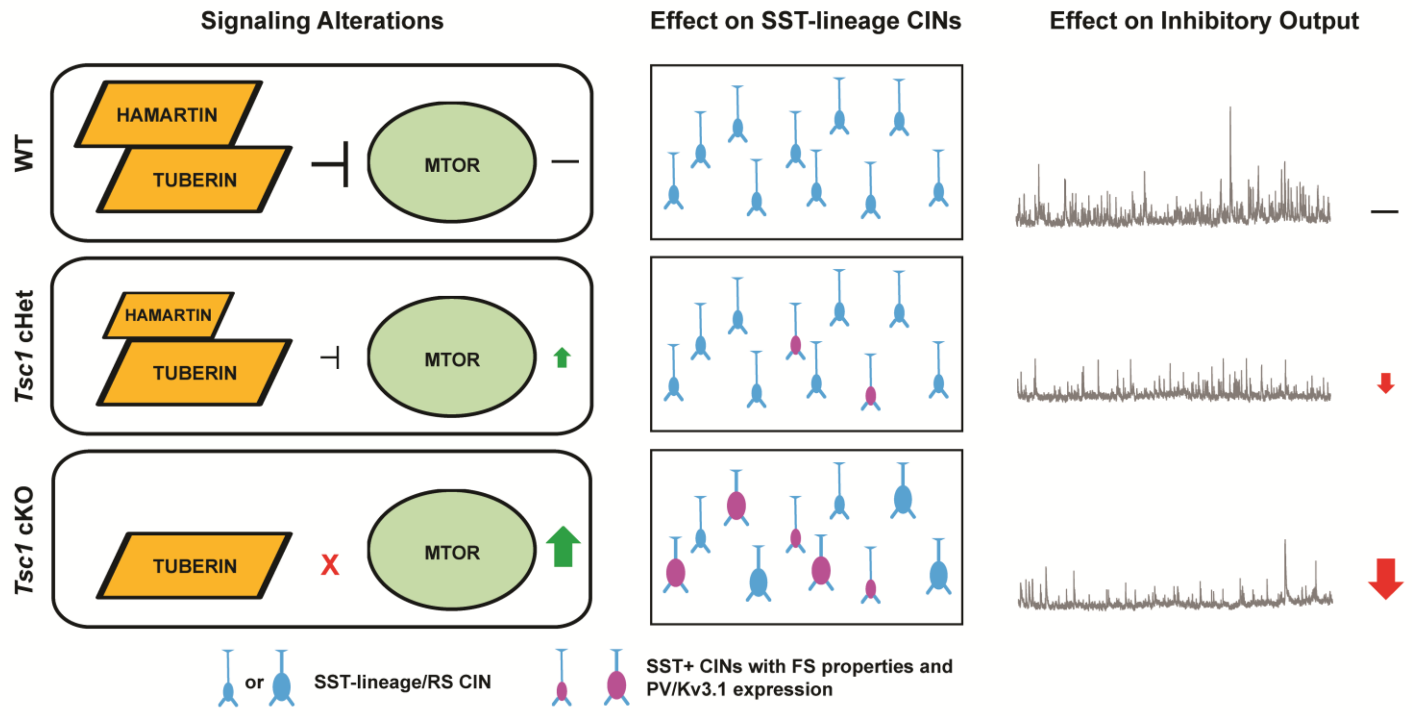


**Model describing the effects of *Tsc1* deletion on cellular properties and synaptic output of post-mitotic SST-lineage CINs in cHets and cKOs.** (Left panels) Loss of either one or two *Tsc1* alleles increases MTOR activity with decreased *Tsc1* dosage (green arrows). (Middle panels) a subset of SST-lineage CINs increase PV expression and exhibit a continuum of physiological properties associated with FS CINs. However, soma size is only grossly increased when both copies of *Tsc1* are depleted. (Right panels) Loss of *Tsc1* in SST-lineages lowers the amount of inhibition in the neocortex, which becomes more impaired as *Tsc1* dosage is decreased (red arrows).
